# Super-resolution expansion microscopy in plant roots

**DOI:** 10.1101/2024.02.21.581330

**Authors:** Michelle Gallei, Sven Truckenbrodt, Caroline Kreuzinger, Syamala Inumella, Vitali Vistunou, Christoph Sommer, Mojtaba R. Tavakoli, Nathalie Agudelo-Dueñas, Jakob Vorlaufer, Wiebke Jahr, Marek Randuch, Alexander Johnson, Eva Benková, Jiří Friml, Johann G. Danzl

**Author notes:** authors contributed equally (Michelle Gallei, Sven Truckenbrodt).

## Abstract

Super-resolution methods enable spatial resolution far better than the optical diffraction limit of about half the wavelength of light (∼200-300 nm) but have yet to attain widespread use in plants, owing in large part to plants’ challenging optical properties. Expansion microscopy improves effective resolution by isotropically increasing physical distances between sample structures while preserving relative spatial arrangements, and clears the sample. However, its application to plants has been hindered by the rigid, mechanically cohesive structure of plant tissues. Here, we report on whole-mount expansion microscopy of *Arabidopsis thaliana* root tissues (PlantEx), achieving 4-fold resolution increase over conventional microscopy, highlighting microtubule cytoskeleton organization and interaction between molecularly defined cellular constituents. By combining PlantEx with STED microscopy, we increase nanoscale resolution further and visualize the complex organization of subcellular organelles from intact tissues by example of the densely packed COPI-coated vesicles associated with the Golgi apparatus and put these into cellular structural context.

## Main

Super-resolution optical microscopy holds tremendous promise for unravelling the molecular organization of plant cells and tissues by increasing our precision in examining biological samples, similar to the transformative power it has had in other areas of biology^1,2^. However, the widespread adoption of super-resolution technologies in plants has been hindered by the unique physiology of plants which make them an optically extremely challenging system. Critically, prominent cell walls and unique organelles like vacuoles produce refractive index variations and pronounced light scattering effects^3^. However, the plant community has started to overcome these challenges to advance our understanding of subcellular plant biology^4,5^, with application of techniques like structured illumination microscopy (SIM)^6–11^, stimulated emission depletion (STED)^12,13^ and REversible Saturable Optical Linear Fluorescence Transitions (RESOLFT)^14^ microscopy, as well as single molecule based methods including photo-activated localization microscopy (PALM)^8,15,16^ and stochastic optical reconstruction microscopy (STORM)^17,18^. As much as these technologies have advanced our ability to visualize plants at improved resolution, some of these provide only an extension of resolution by a factor of two (linear SIM) or below (Airyscan) and hence do not access the sub-100 nm resolution range. Similarly, they are often highly susceptible to aberrations, scattering and fluorescence background, or may require time-consuming generation of new stably expressing fluorescently labelled plant material, or equipment and expertise that is not broadly available.

Expansion microscopy (ExM)^19^ is an elegant strategy to achieve effective resolution better than the diffraction limit, even when imaging with conventional, diffraction-limited microscopes. In ExM, the sample’s structure is imprinted onto a swellable hydrogel that is expanded to increase physical distances between structural elements while conserving their relative arrangements, thus increasing effective resolution. In a crucial step, mechanical cohesiveness of the sample is disrupted by “mechanical homogenization”, either via proteolytic digestion^19^ or denaturation^20^, to abolish counter-forces and enable low-distortion, isotropic, several-fold expansion in each dimension upon wash-out of ions with distilled water. Critically, ExM replaces the need for high-end microscopy equipment and highly trained personnel with a sample-preparation procedure of modest complexity that is readily implemented in any biological laboratory. Furthermore, expanded hydrogels are optically transparent with homogenous refractive index, which mitigates light scattering and aberrations that often hamper other super-resolution approaches in plant samples. ExM also decreases autofluorescence, a notorious adversary in high-resolution plant imaging. Together, this combination of strengths makes expansion microscopy an ideal candidate to aid the adoption and implementation of super-resolution microscopy in the plant biology community. While there have been previous reports of applying hydrogel expansion to plant samples, they were focused on *A. thaliana* ovules/seeds to improve antibody penetration while producing non-uniform 1.2-2.0-fold expansion (likely due to omitting the mechanical homogenization step)^21^, were performed on isolated nuclear organelles^22^ or cells^23^ rather than whole tissue or lacked validation in terms of resolution and distortion^23^.

To overcome the imaging challenges posed by plant samples and to provide an accessible super-resolution imaging methodology for plant tissues, we developed PlantEx (**Fig. 1**), which offers 4-fold resolution increase in three dimensions (3D) based on ExM, using *Arabidopsis thaliana* root as a model system. We evaluated expansion-induced distortions and combined PlantEx with STED microscopy^24–27^ to showcase the road towards molecular-scale optical imaging resolution *in situ* in plant tissues and used comprehensive structural labeling to place super-resolved molecular information into cellular context.

**Figure 1.**
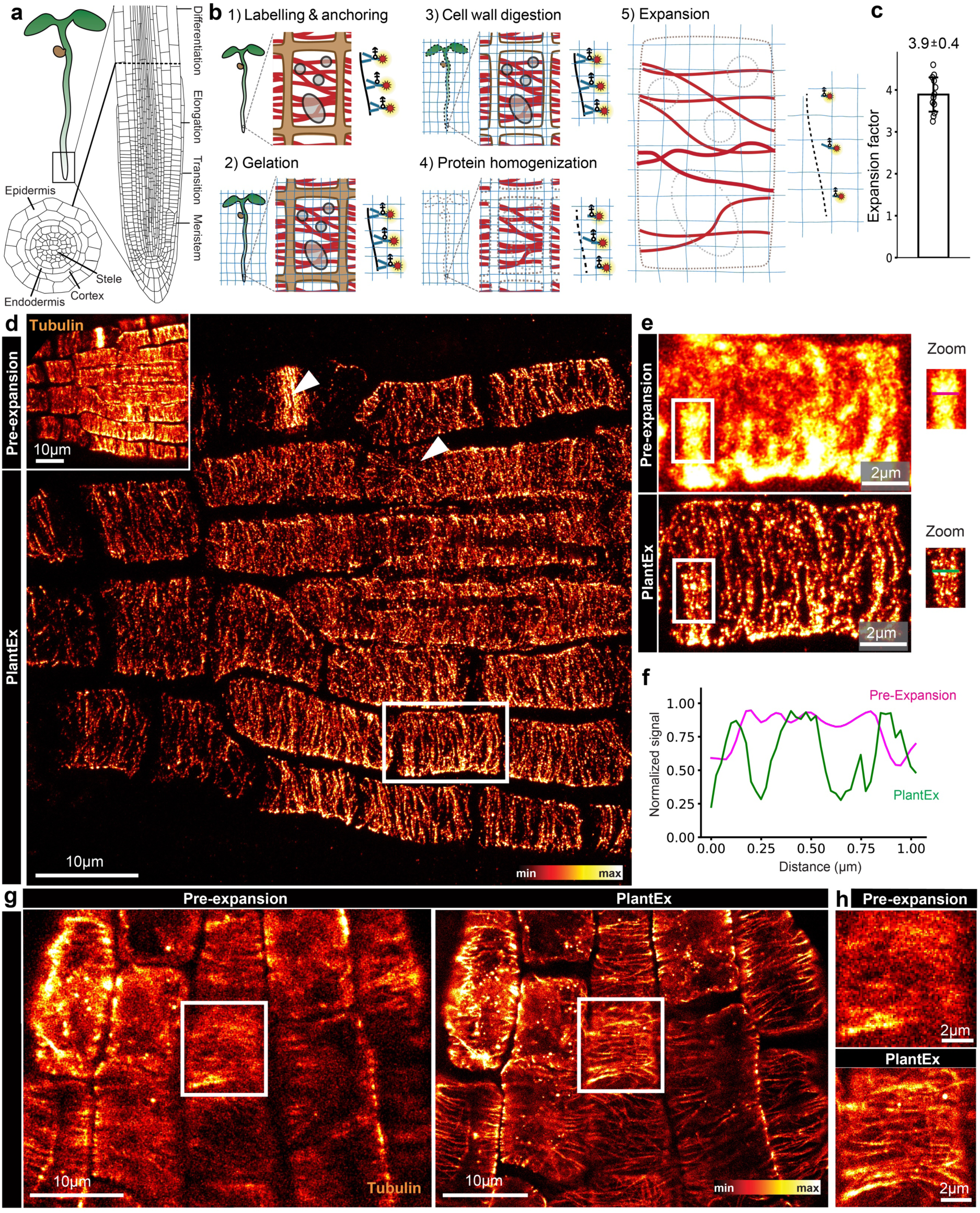
Expansion microscopy of *Arabidopsis thaliana* roots. **a**, *A. thaliana* seedling of ∼4 days age, enlarged root tip, and cross section through root, showing axial zonation and concentric layering of cells. **b**, Schematic of whole-mount expansion microscopy in plants (PlantEx) at whole specimen (left), cellular (middle), and molecular (right) levels. (1) Pre-expansion immunolabeling with fluorophore (red stars) coupled antibodies and addition of anchoring groups. Primary/secondary antibody sandwiches are shown as single antibodies for simplicity. (2) Anchoring groups are copolymerized into an expandable hydrogel. (3,4) A two-step mechanical homogenization procedure disrupts first cell-wall (step 3) and then protein (step 4) cohesiveness by enzymatic digestion. (5) Physical expansion increases effective resolution and renders features distinguishable that cannot be discerned without expansion. Brown: cell walls; red: labelled structures; **c**, Expansion factor for *A. thaliana* root tips, exF = 3.9 ± 0.4 (mean ± s.d., *n*=12 independent specimens). For scaling between pre- and post-expansion data, this exF is used throughout unless otherwise noted. **d**, Confocal images of *A. thaliana* root tip immunolabeled for tubulin before expansion (inset) and with PlantEx, imaged with high-NA objective lens in the lateral root cap. Image scaling between inset and main panel corresponds to expansion by the exF of 3.7 in this measurement. Arrowheads highlight cells of different tubulin organization. Maximum intensity projections of confocal stacks, covering approximately equal axial range in the sample, taking 3.7-fold resolution increase in *z*-direction into account (main panel, 31 slices with 750 nm axial spacing). Scale bars: 10 µm (37 µm physical scale in the expanded sample). Scale bars refer to biological scale, i.e. original tissue size, throughout. Color bar indicates intensity lookup table with saturated pixels indicated in white. **e**, Magnified view of the region boxed in panel (d) for pre-expansion and PlantEx images. Scale bars: 2 µm. *Right*: Additional zoomed views of single confocal image planes before and after expansion in regions marked on the left. Lines indicate position of intensity profiles. **f**, Corresponding line intensity profiles in expanded and non-expanded samples. Distance scale refers to original tissue size. **g**, Single confocal imaging planes in the epidermal cell layer of *A. thaliana* root tip before and after expansion, using improved tubulin immunolabeling with increased labeling density. Scale bars: 10 µm (39 µm physical scale in the expanded sample, exF=3.9). **h**, Magnified view of the cell boxed in g. Images representative of improved tubulin labeling in *n*=4 samples.

## Results

### Establishing expansion microscopy for super resolution imaging of *Arabidopsis* root tissues, cells, and subcellular organelles

Given the success and wide application of ExM to a range of biological systems, we reasoned that ExM would be adoptable to plants if the specific challenges associated with plant anatomy were addressed. In particular, the mechanical cohesiveness of cell walls is expected to hinder expansion if not properly removed, as it is well established that samples with mechanically tough constituents, like e.g. *C. elegans*^28,29^, require specialized protocols for ExM. Even the highly complex but mechanically soft mammalian brain tissue is more challenging to expand homogeneously than monolayer cell cultures and species-dependent differences in composition of thin bacterial cell walls impact expansion^30^. Similarly, tough cuticles of *Drosophila* larvae required a multi-day digestion protocol for expansion^31^. *A. thaliana* cell walls are composed of different polysaccharides, mainly cellulose, hemicellulose, and pectins^32^, and range in thickness from 50 nm to 1 µm, depending on tissue. We established a protocol which overcomes the plant-specific challenges and allows *Arabidopsis* seedlings to be successfully expanded.

We focused the development of PlantEx on the *Arabidopsis thaliana* root organ, as it is a major standard plant species model and the root organ constitutes a popular and fundamental model organ for plant developmental and cell biological studies^33^. *A. thaliana* seedlings (**Fig. 1a**) have a primary root featuring axial zonation^34^ starting at the root tip with the meristematic zone of dividing cells, where also the stem cells reside, and transitioning at the shoot-to-root junction to the hypocotyl-bearing cotyledons. Similarly, root cell files are arranged in concentric layers, encompassing lateral root cap, epidermis, cortex, endodermis, and central cylinder tissue^35^. Each zone has significant physiological roles in development, growth, and regeneration. Intricate spatial organization from tissue to subcellular, organelle, and macromolecular levels mirror important roles in vital processes, such as the gravitropic response and extensive transport with nutrient uptake and synthetic activity.

We based our protocol for PlantEx (**Fig. 1b**) on protein-retention ExM^36,37^ as here, hydrogel embedding and expansion steps are simply added after immunolabeling of specific targets. Importantly, PlantEx can be used with common reporter strategies, including the vast array of plant lines expressing fluorescent protein (FP) fusion proteins. While specialized protocols have been optimized for mild mechanical homogenization to conserve FP fluorescence^37^, we chose to detect FP location and boost signal with fluorophore-conjugated antibodies, which avoided restrictions in designing mechanical homogenization procedures for overcoming cell wall cohesiveness.

Thus, we adopted manual and robotic whole-mount plant immunostaining protocols^38–40,7^, using seedlings aged 3-4 days. These protocols differ from routine mammalian cell labeling protocols as, beyond the standard steps of chemical fixation and membrane permeabilization, plant immunolabeling requires a (mild) cell-wall digestion step permitting antibody access while preserving histoarchitecture. After immunostaining, we incubated whole seedlings with Acryloyl-X solution, providing proteins and specifically fluorophore-bearing antibodies with acrylate anchoring groups that were covalently integrated into the hydrogel upon polymerization. We cast a sodium acrylate/acrylamide/N,N’-methylenebisarcrylamide-based gel designed for 4-fold expansion^19,36^ onto the sample and allowed it to polymerize. We then applied a two-step mechanical homogenization procedure. The first was plant specific and optimized for removing mechanical constraints by cell walls. We found a combination of cell-wall-component specific enzymes driselase, cellulase, macerozyme, pectolyase, and xylanase effective at allowing isotropic expansion of entire root tips. The second homogenization step was akin to homogenization procedures for mammalian tissues to dissolve cohesiveness from protein interactions and relied on proteinase K, a promiscuous proteolytic enzyme. Notably, the plant-specific cell-wall digestion cocktail also displayed promiscuous proteolytic activity (**Suppl. Fig. 1**). It thus complemented proteinase K activity for this purpose, which needs to be taken into account when balancing fluorophore retention against the degree of mechanical homogenization^41^. Immersing the sample/hydrogel hybrid in deionized water resulted in isotropic 3D-expansion while conserving overall sample structure.

To demonstrate how PlantEx can aid the understanding of plant tissue architecture down to the sub-cellular level and to quantify its performance, we focused our super-resolution imaging on the meristematic regions of roots. We immunostained *A. thaliana* seedlings for tubulin and compared confocal pre-expansion and PlantEx images of the same region obtained with high-numerical-aperture objective lenses. Hydrogel expansion conserved cell and tissue morphologies, and, as expected from ∼4-fold expansion, a substantial resolution increase was evident with PlantEx (**Fig. 1d-f**, expansion factor, exF=3.7). We determined the expansion factor as the scale factor between pre- and post-expansion datasets (**Suppl. Fig. 2**) and observed a mean exF=3.9 ± 0.4 (mean ± standard deviation (s.d.)) when evaluating across multiple specimens (**Fig. 1c**, *n*=12 independent specimens). Effective resolution increase equivalent to the exF allowed us to decode the tubulin organizational states in different cells of the tissue at substantially increased detail over pre-expansion images (**Fig. 1d-e**). With PlantEx, line intensity profiles over the same structure showed separated features that coalesced in pre-expansion images (**Fig. 1f**), demonstrating improved resolution with PlantEx. At such high resolution, spatial sampling of the biological structures with fluorophores is a critical factor that may become limiting. In the PlantEx data in **Fig. 1d,e**, the discrete nature of the tubulin staining was apparent, whereas it was concealed in pre-expansion images. Effective labeling density after expansion depends on factors including the efficiency of retention of fluorophore-bearing antibody fragments in the hydrogel^36,37^ and potential loss of fluorophores due to radical generation during polymerization^41^, but equally also on initial antibody penetration into the non-expanded tissue and labeling density achieved before expansion. Therefore, we adopted an immunolabeling protocol specifically optimized for tubulin^7^ and observed improved fluorophore coverage of individual tubulin strands (**Fig. 1g,h**). This showcases that PlantEx, just as other ExM approaches, is a modular technique and improvements in one aspect often directly translate into improved overall outcome. The effect of 3D-resolution enhancement with PlantEx is particularly evident for densely packed structures. When imaging mitotic spindles (**Suppl. Fig. 3**), they appeared as largely homogeneous bright bars before expansion, whereas the organization of tubulin strands could be visualized with PlantEx.

### Characterization of distortions

To determine the fidelity at which cellular structures were expanded with PlantEx, we evaluated the magnitude of expansion-induced distortions. We aligned rigidly scaled pre-expansion and PlantEx images of root samples and calculated displacement vector fields as well as the magnitude of distortions as a function of measurement length (**Fig. 2**). Overlaying overview images of the same tubulin-labeled hydrogel-embedded root taken before and after expansion (**Fig. 2a**) indicated that the expansion procedure maintained overall architecture. We next immunostained for the cell surface markers PIN1 and PIN2 - two auxin transporters located at the cell membrane, expressed in deep root meristem layers and in the outer layers of cortex and epidermis, respectively^42,43^, to evaluate expansion-induced distortions of cell outlines (**Fig. 2b,c**). Here, super-resolution imaging in deep layers of the root was facilitated by optical clearing mediated by hydrogel embedding and expansion, which homogenized refractive index and adjusted it to that of water, and suppressed fluorescence background. We further analyzed distortions at the tissue level (**Fig. 2d**) in the high-resolution data from the tubulin-labeled root in **Fig. 1d**. Taken together, we observed root-mean-squared displacements of a few percent of the measurement length (≲5%) between corresponding pre- and post-expansion images across these measurements (**Fig. 2e**), similar as in previous work on mammalian tissue^44,45^. This demonstrated that expansion of plant tissues was possible at high fidelity with expansion factors equaling other sample types for the chosen gel chemistry. Enzymatic cell wall digestion loosens the rigid polysaccharide structure while maintaining the integrity of plant architecture.

**Figure 2.**
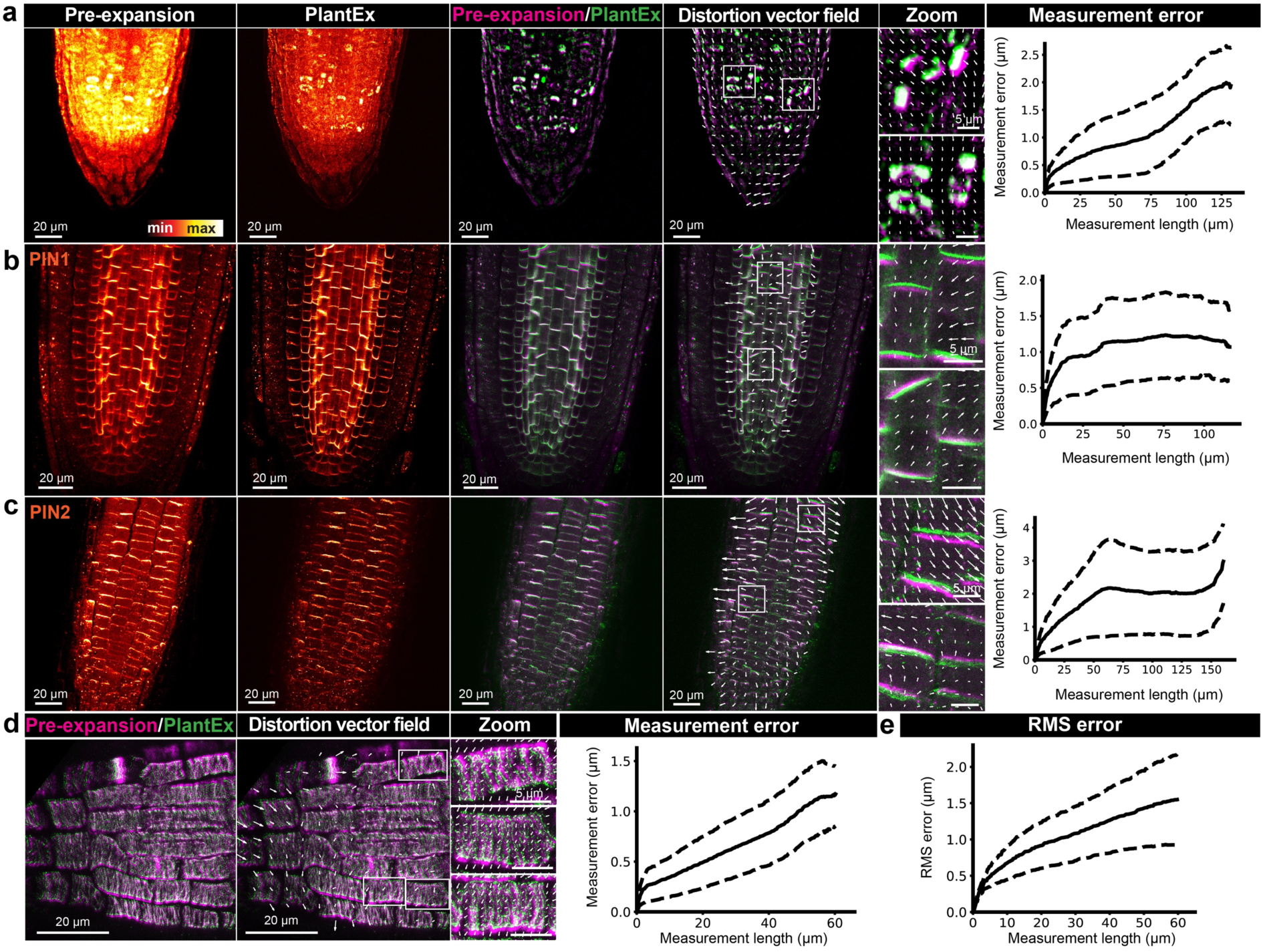
Characterization of distortions. **a**, *Left to right*: Maximum intensity projections of confocal overview image stacks of the same tubulin-labeled *A. thaliana* root tip before and after expansion (imaged with low-NA objective lens). Overlay of scaled and aligned pre-expansion and PlantEx images and distortion vector field with zoomed views characterizing displacements between pre- and post-expansion images. For display purposes, length of distortion vectors was scaled up 2.5-fold in the overview images. To focus distortion analysis on distinct image features, homogeneous components of the signal were removed (for details see Methods and for image alignment see **Suppl. Fig. 2**). Plot of distortions between the pre- and post-expansion images as a function of measurement distance for the same sample. Numerical values for distances and distortions refer to pre-expansion scale throughout. Solid and dashed lines refer to mean and standard deviation, respectively. Scale bars: 20 µm (82 µm physical scale in the expanded sample, exF=4.1). Scale bars, zoomed views: 5 µm. **b,c**, Similar measurements for PIN1 and PIN2 immuno-labelled roots, outlining cell membranes (pre-expansion imaging with 40x, NA1.2 objective lens, PlantEx imaging with 20x, NA0.8 objective). Scale bars: 20 µm (74 µm (PIN1) and 78 µm (PIN2) physical scale in the expanded samples, exF=3.7 and 3.9, respectively). Scale bars, zoomed views: 5 µm. **d**, Overlay of pre- and post-expansion images, distortion vector fields, and distortions as a function of measurement length for the measurement in **Fig. 1d**. Scale bars: 20 µm (74 µm physical scale in the expanded sample, exF=3.7). Scale bars, zoomed views: 5 µm. **e**, Distortions as a function of measurement distance, quantified as root-mean-squared (RMS) error across the *n*=4 samples in panels a-d (see Methods). Solid and dashed lines refer to mean and standard deviation, respectively.

### Dual channel PlantEx

ExM does not require specific fluorophore properties or fluorophore state-control and thus offers a facile route to multi-color super-resolution imaging. We sought to adapt PlantEx for multi-color imaging to further its applications for biological studies exploring nanoscale spatial relationships (**Fig. 3**). We first evaluated the spatial arrangement of Microtubule associated protein 4 (MAP4) and microtubules at extended resolution (**Fig. 3a**), using an established transgenic line^46^ expressing a MAP4-GFP fusion protein. We used GFP as a tag and visualized it with immunolabeling and combined it with immunolabeling for tubulin, clarifying their relative arrangement. We next extended the range of molecular targets and cellular structures by addressing the relationship between components of the Golgi apparatus (GA) and vesicle trafficking machinery^47^ (**Fig. 3b**). The plant GA consists of discontinuous entities dispersed throughout the cytoplasm. COPI-coated vesicles mediate cargo trafficking within the GA and retrograde Golgi-endoplasmic-reticulum trafficking, and are indispensable for plant growth and development^48,49^. To elucidate the spatial interplay between COPI-coated vesicles and the trans-Golgi network (TGN), part of the GA that relays vesicles to various cellular targets, we used an established *A. thaliana* transgenic line^50^ where the TGN was highlighted by expression of GFP fused to the TGN-resident H^+^-ATPase VHA-a1. We detected both the GFP-tag and endogenous Sec21, the γ-subunit of COPI-coated vesicles, via immunostaining. Without expansion, using a high-NA objective lens for confocal imaging, these markers were largely overlapping and it was not possible to determine their spatial relationship. In contrast, in PlantEx, the arrangement of the individual COPI-coated vesicles around the TGN perimeter was readily discernible (**Suppl. Movies 1,2**). This agrees with previous observations of COPI-vesicles budding from and trafficking between Golgi cisternae^48^. Also here, the signal from immunolabeled cellular structures stood out more clearly against the cellular background in PlantEx. With dual channel imaging, PlantEx thus facilitated detailed studies of cellular organization and relative molecular arrangements in plant samples.

**Figure 3.**
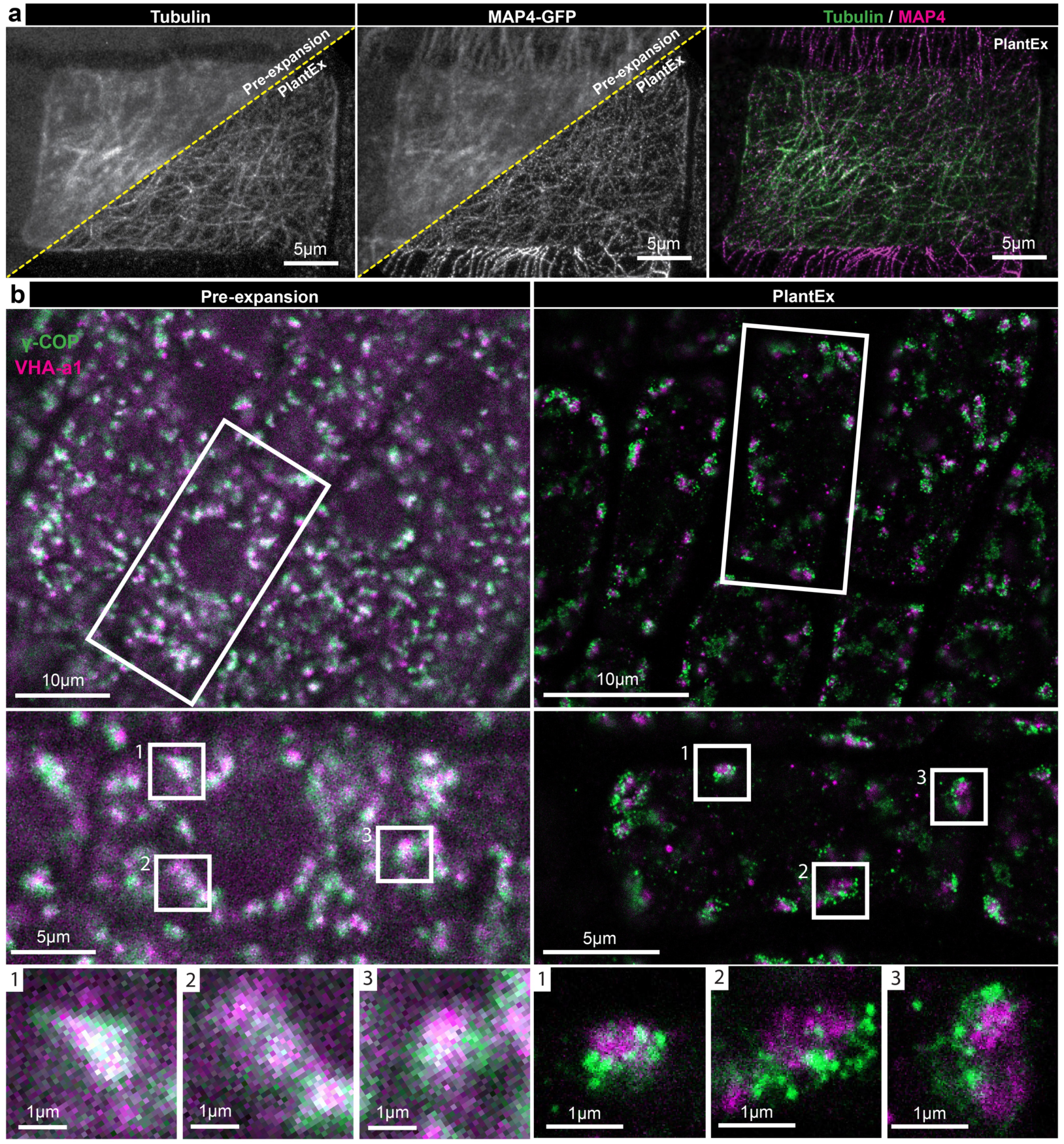
Dual color PlantEx. **a**, Dual-color PlantEx measurement of tubulin and MAP4 in root tissue of an *A. thaliana* line expressing a MAP4-GFP fusion protein. Tubulin and GFP were visualized via immunolabeling. *Left*: Confocal PlantEx image of tubulin in a root cell, with the corresponding pre-expansion confocal image in the upper left corner. *Middle*: MAP4 channel. *Right*: Overlay of tubulin and MAP4 signal. In the specimen imaged here, intensity of pre-expansion immunolabeling was variable between cells. Scale bar: 5 µm. **b**, Dual-color PlantEx measurements of COPI-coated vesicles and trans-Golgi network. *Left*: Pre-expansion confocal images of lateral root cap tissue immunostained for GFP expressed as fusion protein with a trans-Golgi network marker (VHA-a1, magenta) and for the γ-subunit of COPI-coated vesicles (γ-COP, sec21, green). Cell boundaries can be discerned as regions devoid of stained structures. *Right*: Confocal image of a similar region in the same specimen after applying PlantEx. Resolution and signal-to-background ratio were increased. Scale bars: 10 µm (39 µm physical scale in the expanded sample, exF=3.9). *Centre, bottom*: Magnified views of regions indicated by white boxes at cellular and subcellular scales. Examples with roughly corresponding structures were selected from pre-expansion and PlantEx images. No attempt was made to find the exactly corresponding sample regions in the pre-expansion and PlantEx tissue volumes. Color maps are linear with saturation of brightest pixels. Scale bars: 5 µm (center) and 1 µm (bottom).

### Combining PlantEx with existing super-resolution techniques

While PlantEx markedly increases the resolution to visualize *planta* structures, it can be advantageously combined with advanced optical readout modalities (**Fig. 4**). For example, PlantEx with diffraction-limited confocal readout failed to resolve individual COPI coated vesicles in densely packed regions of the cell (**Fig 4a**). Thus, we sought to combine PlantEx with the super-resolution imaging technique STED^24,25,51^ microscopy to further improve the capability to resolve such sub-diffraction organelles. While STED microscopy has been applied to plants^12,13^ it is usually difficult to perform in thick and opaque samples such as plant roots and thus often fails to produce equivalent performance as in other biological samples (**Suppl. Fig. 4**), with STED performance being sensitive to aberrations and scattering. However, as PlantEx provides optical clearing and refractive index matching, we reasoned that it would facilitate STED readout. Using a water immersion objective lens, we applied STED microscopy to expanded hydrogels^26,27^ of *Arabidopsis* seedlings immunolabeled for COPI-coated vesicles with a high-performance STED dye compatible with radical-based hydrogel polymerization (**Fig. 4a**). This PlantEX-STED modality allowed us to capitalize on the combined resolution improvements offered by the individual PlantEx and STED methodologies. Dialing in STED resolution increase in the focal plane (*xy*-direction) in addition to PlantEx resolved individual vesicles that coalesced with conventional confocal readout (**Fig. 4a-c**). The apparent diameter of COPI puncta in such PlantEX-STED images (**Fig. 4d, Suppl. Fig. 6**) was 52 nm (median full-width-at-half-maximum (FWHM), native tissue scale, lower and upper quartiles: 43nm and 63nm, 74 vesicles recorded across *n*=3 specimens). This may be taken as a proxy for the effectively achieved resolution in this dual super-resolution mode, with the caveat that COPI coated vesicles have a size of tens of nm (∼45 nm) themselves^48,52^. Similarly, irrespective of the applied super-resolution readout technology, immunolabeling for an organelle protein component does not necessarily trace out its nanoscale shape due to a potentially limited coverage of the vesicle outline by the finite number of protein targets present, finite epitope coverage, and displacement of fluorophores from the target structure due to non-negligible size of antibodies used for labeling (i.e. linkage error between biological target and fluorophore).

**Figure 4.**
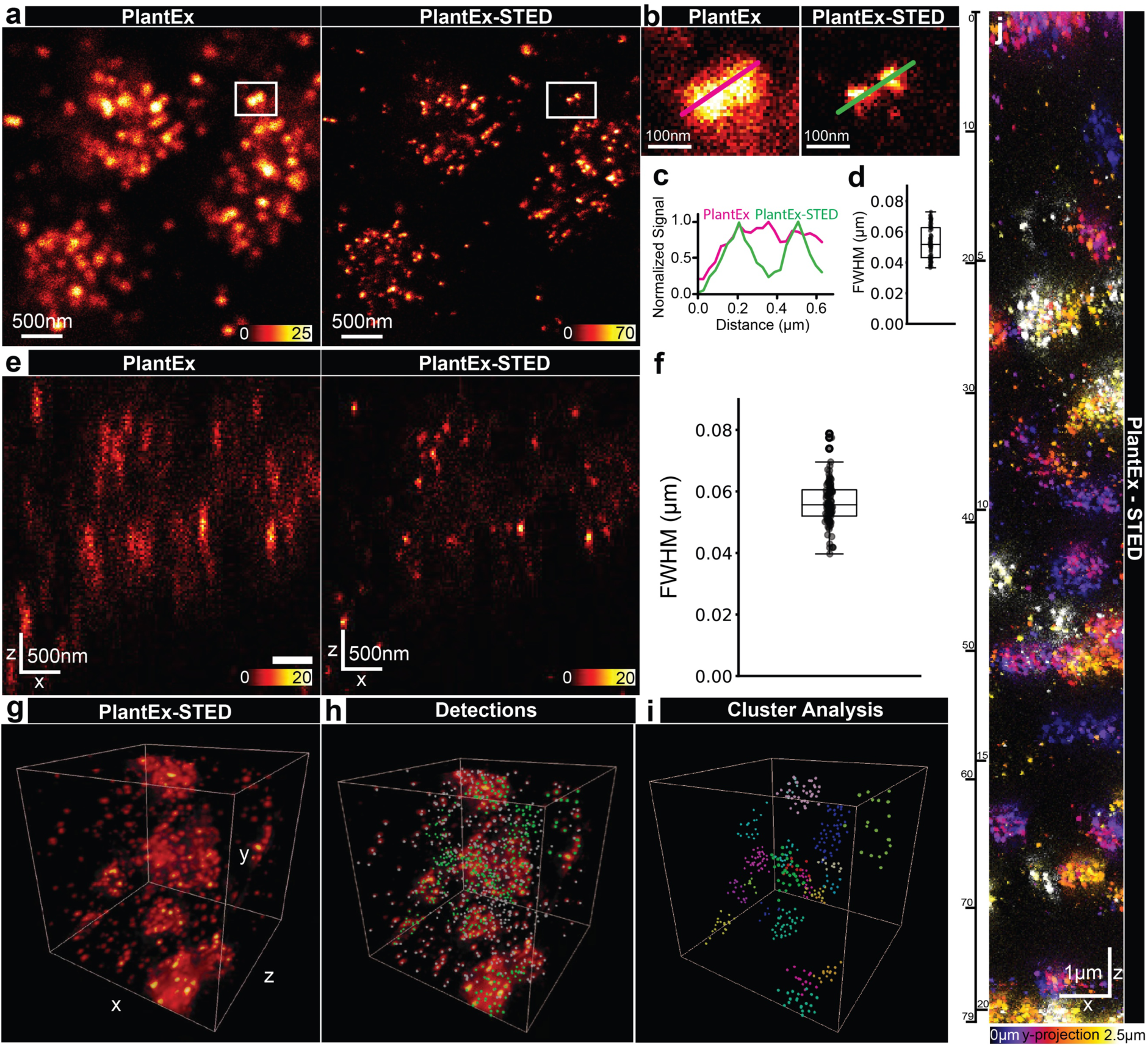
PlantEx with STED microscopy readout. **a**, *Left*: COPI-coated vesicles in an expanded *A. thaliana* whole mount root sample labeled for sec21 (PlantEx), imaged with confocal microscopy. *Right*: Same region imaged with STED microscopy (PlantEx-STED), using a *xy*-STED light pattern (“doughnut” pattern, lateral resolution increase). Scale bar: 500 nm (1.95 µm physical scale in the expanded sample). Numbers in intensity lookup bar represent single-photon detection events. Color map is saturated at the brightest pixels (white). Images representative of STED experiments in *n*=4 specimens. **b**, Magnified views as indicated by boxed regions in a. Lines indicate positions of intensity profiles. **c**, Corresponding line intensity profiles. Distance scale refers to original tissue size. **d**, Full-width-at-half-maximum (FWHM) of 2D-Gaussian fits to γ-COP (sec21) positive puncta in PlantEx-STED with lateral resolution enhancement (median value: 52nm, lower and upper quartiles: 43 nm and 63 nm, 74 puncta across 3 specimens). **e**, *Left*: Axial confocal scan (*xz*-direction) in the same PlantEx sample. *Right*: Axial scan of the same region with STED light pattern (“*z*-STED pattern”) increasing resolution predominantly along the optical axis (*z*-direction). Scale bar: 500 nm (1.95 µm physical scale in the expanded sample). **f**, Full-width-at-half-maximum (FWHM) of 3D-Gaussian fits to γ-COP positive puncta in volumetric PlantEx-STED with *z*-STED pattern, recorded across *n*=3 specimens (median: 56 nm, lower and upper quartiles: 52 nm and 61 nm, 112 puncta). **g**, 3D-rendering of PlantEx-STED imaging volume with labeling for COPI-coated vesicles (γ-COP) imaged at near-isotropic resolution (*z*-STED pattern). Fly-through: **Suppl. Movie 3**. Imaging volume: 3.8 x 3.8 x 3.8 µm^3^ (15 x 15 x 15 µm^3^ physical scale in the expanded sample). **h**, Visualization of γ-COP positive puncta, detected as local intensity maxima in the same volume. **i**, Color coding after applying a clustering algorithm. Puncta not assigned to clusters are indicated in grey (**Suppl. Movie 7**). **j**, PlantEx-STED imaging with extended axial range. COPI-coated vesicles imaged at near-isotropic resolution (*z*-STED pattern) in a tissue column of 79 µm imaging range in *z*-direction, corresponding to 20.25 µm in native tissue. Side view (*xz*-direction), the third dimension (*y*) is color-coded. Scale bar: 1 µm (3.9 µm physical scale in the expanded sample). Representative of volumetric STED imaging in *n*=3 samples.

Given the anisotropic shape of the confocal point-spread function (PSF), we expected that the pivotal factor for discerning vesicles in 3D would be increasing resolution in the axial (*z-*)direction. We thus applied a STED light pattern for resolution increase predominantly in *z*-direction (*z*-STED, **Fig. 4e**). Dialing in near-isotropic diffraction-unlimited resolution and employing RESCue (Reduction of State transition Cycles)-STED^53^ to mitigate photobleaching allowed volumetric PlantEx-STED imaging (**Fig. 4g, Suppl. Fig. 5, Suppl. Movies 3-6**). This was compatible with detection of COPI-coated vesicles by a straightforward peak-finding algorithm, whereas they would largely coalesce in conventional diffraction-limited images or PlantEx alone. Automated detection allowed for visualization of their 3D spatial arrangement in tissue and we used a clustering algorithm to color code detections and highlight regions of high vesicle abundance (**Fig. 4h,i**, **Suppl. Movies 7-10**). We also quantified the apparent size of the COPI-puncta in volumetric, near-isotropically resolved PlantEx-STED datasets with 3D-Gaussian fits, resulting in a median FWHM of 56 nm (native tissue scale, lower and upper quartiles: 52 nm and 61 nm,112 vesicles recorded across *n*=3 specimens) (**Fig. 4f, Suppl. Fig. 6**).

In tissue imaging, it is typically challenging to achieve super-resolution over an extended axial range, due to depth-dependent aberrations (e.g. spherical), scattering, and photobleaching. Hydrogel-based refractive index homogenization in combination with a high-NA water-immersion objective lens largely solves these issues. Together with RESCue-STED, this allowed us to image the 3D-distribution of COPI-coated vesicles in a hydrogel column of ∼80 µm axial extent (**Fig. 4j**), corresponding to ∼20 µm in the native tissue, within a single measurement. Together, this demonstrated that PlantEx can be advantageously combined with advanced optical readout modalities, as exemplified here with STED microscopy to improve overall achievable resolution.

### Comprehensive visualization of tissue architecture in PlantEx

We were next interested in providing a means to comprehensively visualize tissue architecture at super-resolved subcellular scale with PlantEx. We hence applied fluorophores bearing an N-hydroxysuccinimide (NHS) ester to PlantEx samples (**Fig. 5, Suppl. Fig. 7**), inspired by visualization of cell and tissue architecture in mammalian cells and tissues^54–57^ through indiscriminate (“pan”) protein-density staining. These covalently reacted with (primary) amines abundant on cellular proteins and thus mapped cellular structure comprehensively. Such labeling clearly outlined cell shapes and we used it to place specific molecular signals into the context of subcellular architecture at resolution 4-fold increased by PlantEx (**Fig. 2b-e**). For this, we imaged the structural channel with confocal microscopy and visualized the TGN and COPI-coated vesicles at further augmented resolution with STED microscopy (**Fig. 2b-c**). Next, we applied NHS-ester modified STED-compatible fluorophores to expanded root samples, thus enabling further resolution increase with STED also in the structural channel (**Fig. 2d**). Protein-density labeling in PlantEx also provided cellular context in volumetric datasets, when we imaged COPI-coated vesicles in PlantEx-STED mode at near isotropic resolution (using a *z*-STED pattern) and read out the pan channel with confocal microscopy (**Fig. 5e**).

**Figure 5.**
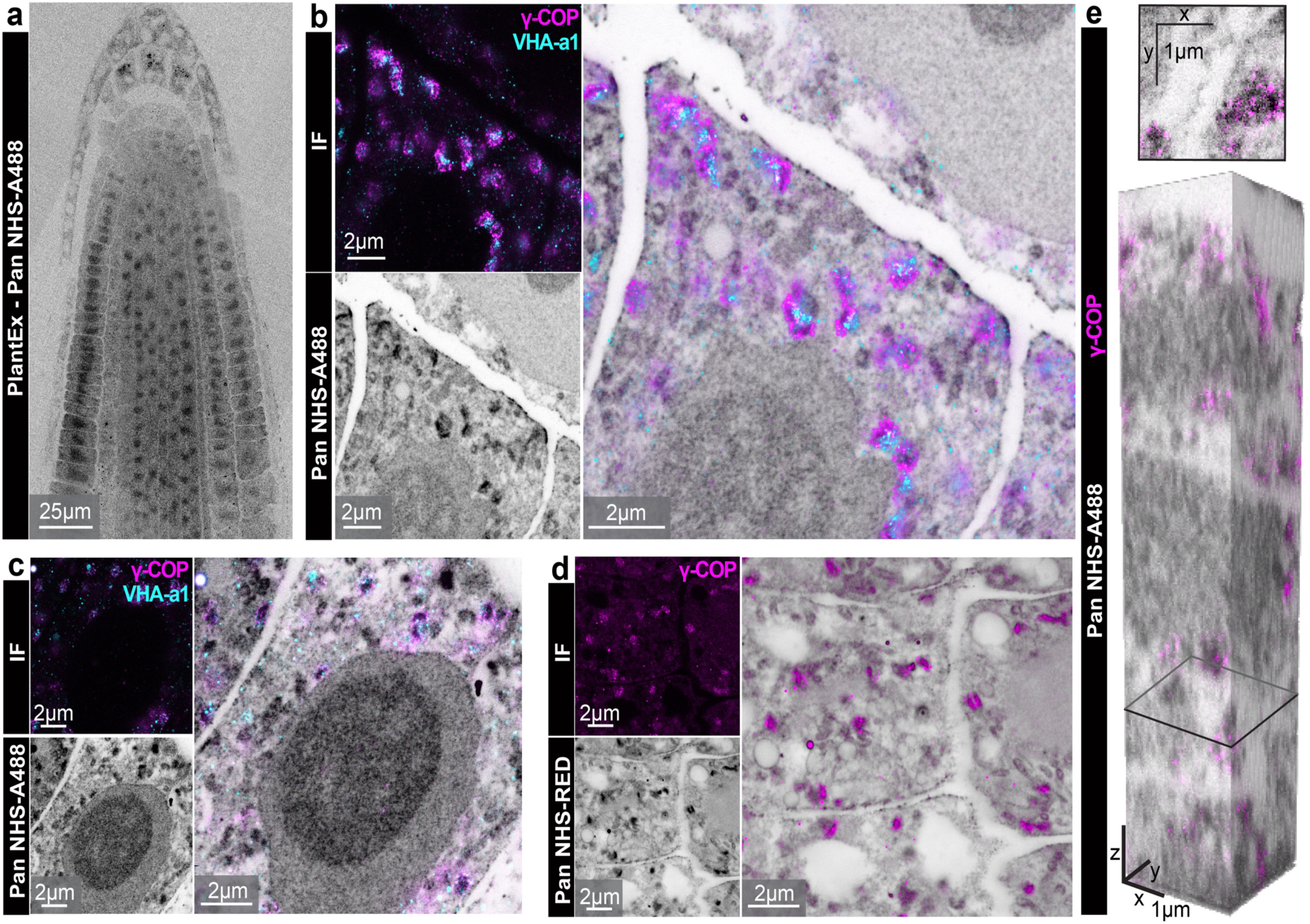
PlantEx with comprehensive visualization of tissue architecture. **a**, Spinning-disk confocal overview image of expanded *A. thaliana* whole mount root sample with “pan”-labeling using an NHS-modified fluorophore (grayscale, NHS-A488: N-Hydroxysuccinimide-Alexa 488) to map protein density. Grayscale color map is inverted (black represents high protein density, non-inverted images: **Suppl. Fig. 7**). Scale bar: 25 µm (97.5 µm physical scale in the expanded sample). **b**, Trans-Golgi network and COPI-coated vesicles in cellular context. *Top left*: PlantEx-STED image of VHA-a1 (cyan, visualized by immunolabeling for GFP-tag, *xy*-STED) and γ-COP (immunolabeling for sec21, magenta, *xy*-STED) in whole-mount *A. thaliana* root PlantEx specimen. *Bottom left*: Same region visualizing cellular architecture with pan-protein label (confocal, grayscale, color map inverted, NHS-A488). High intensity in pan-channel at TGN/vesicle locations partially reflect increased protein density from antibodies applied before NHS-fluorophore labeling. *Right*: Overlay. Scale bars: 2 µm (7.8 µm physical scale in the expanded sample). **c**, Similar measurement in different region (γ-COP with *xy*-STED, VHA-a1 confocal). Scale bars: 2 µm. **d**, PlantEx of cellular architecture in whole mount *A. thaliana* root sample with pan-protein labeling, read out with STED microscopy (grayscale, *xy*-STED, NHS-RED: NHS-Star Red, colormap inverted), with COPI coated vesicles (γ-COP, magenta, confocal). Scale bars: 2 µm. For the overlay, background counts in the immunolabeling channel were subtracted for visual clarity. **e**, Volumetric PlantEx-STED imaging with cellular context. COPI-coated vesicles (γ-COP, magenta, *z*-STED) in cellular context of pan-protein labeling in *A. thaliana* PlantEx root sample (confocal, grayscale, NHS-A488, colormap inverted). *Top*: Cross section at the position indicated in the bottom panel. *Bottom*: 3D-rendering of volume (2.6 x 2.6 x 12.8 µm^3^ native scale). Scale bars: 1 µm.

We also found that indiscriminate protein labeling was practically useful to locate the otherwise transparent PlantEx samples in 3D within the hydrogel for imaging, also when using a low-NA objective lens with moderate photon collection efficiency. More importantly, comprehensive structural labeling provided a general and straightforward way of assessing tissue architecture and preservation in expanded plant samples (**Fig. 5a**). As plant seedlings are delicate specimens, we found this a convenient way to rapidly gauge whether a particular specimen had been damaged during manual handling for fixation, labeling, or hydrogel embedding and it allowed for an instantaneous assessment of homogeneity of expansion independent of immunolabeling. Taken together, pan protein labeling substantially extends the information content in PlantEx imaging.

## Discussion

Here we developed expansion microscopy for plant tissues, using the intact *Arabidopsis thaliana* root as model system, thus overcoming the notoriously unfavorable optical properties of plants for super-resolution imaging. PlantEx offers 4-fold increase in effective resolution over the chosen readout modality in all three spatial directions for multiple color channels, and preserves *planta* structures with high fidelity. This provides effective super-resolution also with widely available conventional diffraction-limited microscopes, such as confocal systems. It thus opens up the prospect of nanoscopic imaging studies for plant biology laboratories not specialized in, or lacking access to, classical super-resolution imaging. We showcased how PlantEx may facilitate biological discovery, visualizing multiple molecular targets including cytoskeletal organization and nanoscale arrangement of organelles. Notably, PlantEx is compatible with existing well-characterized fluorescent protein expressing marker lines by immunolabeling for the fluorescent protein tag. We also demonstrated how PlantEx can be further enhanced choosing advanced optical readout modalities according to the application requirements. We combined it with volumetric STED super-resolution imaging in the intact tissue to resolve individual COPI-coated vesicles in dense clusters associated with the Golgi apparatus. We also used this system to highlight that the improved resolution and signal-to-noise ratio offered by PlantEx facilitate analysis of biological specimens with automated approaches.

As in any expansion technology, resolution increase in PlantEx is equivalent to the expansion factor (4x, ∼65 nm lateral resolution at 490 nm excitation with confocal readout using an NA=1.2 objective lens). Hydrogel embedding and expansion also optically clear the sample, by removing cellular constituents such as lipids, cell wall components and chlorophyll, and by homogenizing and adjusting the refractive index to that of water. Endogenous fluorophores likely fail to be anchored or may be destroyed during polymerization and homogenization. The rapid nature of PlantEx sample preparation offers advantages over existing time-consuming plant clearing methods^58^. Clearing and index matching benefit optically demanding readout modalities. For instance, the quality of the intensity minimum of a *z*-STED pattern for axial resolution increase^25^ is highly sensitive to spherical aberrations and scattering, posing a challenge in native plant samples whereas this is largely abolished when using water immersion objectives with PlantEx hydrogels, even when imaging whole-mount samples as we did here. It stands to reason that PlantEx can advantageously be combined with further optical imaging modalities depending on measurement requirements, such as lightsheet microscopy.

The currently achieved 4-fold resolution increase compares favorably against the more moderate gains in resolution with Airyscan or linear SIM, while these modalities bear the advantage of being also applicable to living samples. Also RESOLFT^14^ is applicable to living specimens but requires specialized equipment and engineering of transgenic lines expressing fluorescent proteins with specific (photoswitching) properties. Optical super-resolution techniques have been applied to plant samples to achieve high resolution, down to e.g. 40 nm in the case of STORM^18^. However, optical aberrations, scattering, and fluorescence background do render these challenging for tissue imaging, especially for accessing deeper portions of the tissue. Through combination with STED readout, it was straightforward to achieve resolution increase in PlantEX beyond the 4-fold expansion. Incorporating alternative hydrogel chemistries^45,57,59,60^, may in the future provide a route towards further resolution improvements using standard microscopes.

Beyond the obvious restriction of ExM to fixed specimens, a series of considerations apply when adopting the technology. Common to all super-resolution approaches, the technology visualizes fluorophores rather than the actual biological target molecules. Hence, biological structures need to be sampled by fluorophores at sufficient labeling density for adequate visualization. While the discrete spatial sampling by fluorophores is typically concealed at diffraction-limited resolution, it often becomes apparent at the tens of nm resolutions achieved with PlantEx and even more so when resolution is further increased in PlantEx STED. The overall imaging outcome thus crucially depends on the quality of the initial antibody staining and we showed how an improved immunolabeling protocol enhanced visualization of microtubules in PlantEx. In our protocol, hydrogel embedding and expansion ensued after immunolabeling. It is hence only applicable to targets where labeling can be performed in the native tissue. Despite the initial, mild cell wall digestion step, antibody penetration into the tissue may vary across specimens and may result in labeling restricted to superficial cell layers. Common to all expansion approaches with pre-expansion immunolabeling, only labels retained in the hydrogel can be detected post expansion. Well-known tradeoffs exist in protein-retention ExM between the degree of mechanical homogenization and retention of fluorophore-bearing antibody fragments after proteolytic treatment^41^. Stronger digestion increases fluorophore loss and we point out that certain fluorophores, notably cyanine dyes, are not compatible with the radical chemistry during hydrogel formation. Specific to PlantEx, the plant-specific cell-wall digestion cocktail also displays proteolytic activity, which should be taken into account when optimizing mechanical homogenization including proteinase K digestion. Specific optimization may indeed be required, as the cocktail includes natural extracts whose activity may vary between batches. Another caveat common to approaches involving (pre-expansion) immunolabeling is a displacement of fluorophores from target structures, as fluorescence imaging visualizes fluorophores rather than the molecules or structures of interest themselves. Such linkage error may reach ∼20 nm with the secondary immunostainings applied here, as antibodies are large molecules of ∼150 kDa and ∼10 nm size. Further, the spatial arrangement and orientation of antibodies with respect to the target epitopes is not known. A low copy number of target proteins on individual cellular organelles or incomplete epitope coverage may impede outlining the precise shape of an organelle. Consequently, measurements based on protein antibody labeling of vesicle proteins are useful to put bounds on resolution rather than for determining the precise organelle size. There is potential for improving these various factors by development of approaches using smaller probes (e.g., nanobodies), post-expansion labeling^20^, or optimized signal retention strategies^61^. For quantitative biological measurements, validation strategies for each cellular target must be adapted to the biological question posed, including homogeneity of expansion of the target structure.

Beyond the obvious benefit of visualizing tissue architecture comprehensively and providing super-resolved subcellular context, we found protein-density mapping (“pan-protein” labeling) a useful way to assess structural preservation for individual samples and to locate the plant root in the expanded hydrogel, as the cleared seedlings produce hardly any non-fluorescent contrast and immunolabelings can be faint.

Our protocol was developed using the *A. thaliana* root organ, thus enabling cell biological and developmental studies in this important model system. We expect that this will also spark adaptation to other species. This will require tuning of the labeling and mechanical homogenization procedures, as different species are expected to feature different molecular and mechanical characteristics, in particular regarding cell wall composition. Similarly, expansion of different organs will require dedicated protocols. Both aspects are akin to the application of expansion microscopy to animal samples, where a wide variety of protocols have been developed to cater for differences according to species or tissue imaged.

Overall, PlantEx enables accessible super-resolution imaging to facilitate novel cell biological studies that examine subcellular entities and their mutual spatial relationships. Furthermore, PlantEx will enable the decoding of complex 3D-topologies of cellular organelles and the spatial interplay with interaction partners, giving access to cellular nanoscale architecture in a molecularly informed way.

## Acknowledgements

We gratefully acknowledge support by the Scientific Service Units at ISTA, including the Imaging and Optics and Lab Support facilities and the mechanical workshop and Library. We thank Philipp Velicky for STED microscope alignment.

This project has received funding from the Austrian Science Fund (FWF): I 3630-B25 (J.G.D) and the European Research Council (ERC) under the European Union’s Horizon 2020 research and innovation programme (grant agreement No 742985, J.F.). It has also received funding from the European Union’s Horizon 2020 research and innovation programme under the Marie Skłodowska-Curie Grant Agreement No. 665385. S.T. has received funding as an ISTplus Fellow from the European Union’s Horizon 2020 Research and Innovation Programme under Marie Skłodowska-Curie grant agreement no. 754411 and from an EMBO Long-Term Fellowship (grant number ALTF 679-2018). It has further received funding from the Austrian Science Fund (FWF) grant DK W1232 (M.T, N.A-D., J.G.D). W.J. received funding via a Human Frontier Science Program postdoctoral fellowship LT000557/2018.

The funders had no role in study design, data collection and analysis, decision to publish or preparation of the manuscript.

## Author contributions

M.G., S.T., C.K. and S.I. performed experiments. M.G., S.T., C.K., V.V. and C.S. analyzed data. M.T., N.A.-D., J.V., W.J., M. R. and A.J. supported experiments and imaging. E.B. and J.F. provided samples and advised on labeling and plant biology. J.G.D. conceived and supervised the study. M.G., S.T., C.K., V.V., and A.J. prepared figures. M.G., S.T., A.J. and J.G.D. wrote the manuscript with input from all authors.

## METHODS

### Plant material and growth conditions

*Arabidopsis thaliana* ecotype Col-0 was used for tubulin staining. For imaging of COPI-coated vesicles and trans-Golgi network (TGN), a VHA-a1-GFP expressing line^50^ with Col-0 background was used. For visualization of MAP4, a MAP4-GFP line^46^ with Col-0 background was used. Seeds were surface-sterilized with chlorine gas (generated from 39 ml 12 % sodium hypochlorite-based bleach, 61 ml H_2_O, and 4.5 ml 37% HCl) overnight or with a 15 minute 70% ethanol wash followed by a 15 minutes 100% ethanol wash and then air-dried under sterile conditions. Following a 48-72 hour 4°C stratification, seedlings were germinated and grown on ½ MS (Murashige-Skoog) medium (1% D+ Saccharose (Sigma 84100), 0.23% Murashige-Skoog powder (Duchefa M0221.0050), 0.05% MES (2-(N-morpholino)-ethanesulfonic acid) (Duchefa M1503.0100), 0.8% Agar (Duchefa P1001.1000), with pH set to 5.6 with KOH prior to autoclaving) for 3-5 days in a growth room at 21°C and long-day conditions (16 hours light, 8 hours dark).

## Fixation and immunostaining

Fixation was done with paraformaldehyde and immunostaining was either performed manually or with a pipetting robot. Labeling protocols included a mild first cell wall digestion step to allow for antibody access but maintain tissue integrity. Thorough cell wall digestion for PlantEx was performed after anchoring and hydrogel polymerization.

*Tubulin labeling for ExM.* For tubulin staining, we followed a previously published fixation and labeling protocol^40^. Seedlings of 3 to 4 days of age were fixed in a 12 well plate with 2% paraformaldehyde in 1x MTSB (microtubule stabilization buffer) supplemented with 0.1 % Triton X-100 (Sigma-Aldrich, T8787). 2x MTSB stock solution was: 15 g PIPES (piperazine-N,N′-bis(2-ethanesulfonic acid, Sigma-Aldrich, P1851)), 1.90 g EGTA (ethylene glycol-bis(β-aminoethyl ether)-N,N,N′,N′-tetraacetic acid, Sigma-Aldrich, E3889), 1.22 g MgSO_4_·7 H_2_O (Merck Chemicals, 230391) and 2.5 g KOH (Merck Chemicals, 105033) in 500 ml water, adjusted with KOH to pH 7.0. Samples were placed in a vacuum desiccator and suction was applied for 5 minutes, pressure brought back to ambient values, and suction applied again for 5 minutes to facilitate penetration of fixation solution. Fixation proceeded for additional 40 minutes at 37 °C and samples were washed with water and placed overnight in water at 4 °C. Next, seedlings were incubated for 10 minutes in 800 μl methanol at 65 °C, which solubilizes the plant cuticle and further components, such as chlorophyll. We then added 200 μl of Millipore water every 2 minutes until the final alcohol concentration reached ∼20% and washed seedlings with water. Cell wall digestion for antibody access was performed with a solution containing 0.2% driselase (Sigma-Aldrich, D8037) and 0.15% macerozyme R-10 (Yakult Pharmaceutical Industry Co., Ltd.) in 2 mM MES, pH 5.0 for 40 minutes at 37 °C. After washing in MTSB, membranes were permeabilized with 3% IGEPAL CA-630 (Sigma-Aldrich, I8896), 10% DMSO (Sigma-Aldrich, W387520) in 1x MTSB for 20 minutes at 37 °C. After 4 washes in 1x MTSB, samples were blocked in 1x MTSB supplemented with 2% BSA (Bovine Serum Albumin, Sigma-Aldrich, A2153) for 30 minutes and incubated first with primary antibody (anti-Tubulin, Santa Cruz Biotechnology, SACSC-53030, monoclonal, IgG2a, 1:100, in blocking solution), washed, and then with secondary antibody (Alexa Fluor 488, Goat anti-Rat IgG (H+L) cross-adsorbed secondary antibody, Invitrogen, A 11006, 1:100, in blocking solution) for 2 hours each and washed again. If anchoring was not done on the same day, samples were stored in storage solution containing 90% Glycerol (VWR, ICNA0219399691), 10% PBS, DABCO (25 mg/ml, 1,4-Diazabicyclo[2.2.2]octane, Sigma Aldrich, D27802), with pH adjusted to 9.

*Optimized Tubulin labeling.* Manual fixation and immunolabelling were performed according to a previously published protocol^7^, with minor modifications. 3 to 4 day old *A. thaliana* seedlings were treated with 10nM Taxol for 30 minutes and then were fixed in 1.5% formaldehyde, 0.1% glutaraldehyde with 0.01% Triton X-100 in MTSB for 1 hour at room temperature. Samples were gently transferred onto a chambered slide and incubated in room temperature for 30 minutes in a cell-wall digesting solution of 2.5% Cellulase Onozuka R10 (Yakult Pharmaceutical Industry Co.), 1% Macerozyme R10 (Yakult Pharmaceutical Industry Co.), 1% Meicelase (Desert Biologicals), 0.1% Pectolyase (Duchefa, P8004) in MTSB. Samples were washed 3x with MTSB and 3 times with PBS pH7.4 for 5 minutes each at room temperature. Since glutaraldehyde was used for fixation, the seedlings were then incubated for 15 minutes in 0.1% sodium borohydride in PBS. Following this aldehyde reduction step, the samples were permeabilized by 15 minutes incubation in 10% DMSO, 2% IGEPAL CA-630 (Sigma-Aldrich, I8896) and 0.01% Triton X-100 in PBS. Samples were then washed in PBS and were incubated overnight in a blocking solution consisting of 3% BSA (Bovine Serum Albumin, Sigma-Aldrich, A2153) and 0.5% BSA-c(polyacetylated BSA, Aurion) in PBS. Seedlings were first incubated overnight at room temperature in primary antibody solution (anti-α tubulin, mouse monoclonal, Sigma Aldrich, T6074, 1:100 and anti-β tubulin, mouse monoclonal, Sigma Aldrich, T5293, 1:100) followed by 6 washes in PBS for 10 minutes each. Optionally, for MAP4-GFP detection, additional labeling was performed with rabbit anti-GFP (Agrisera, AS152987) and using rat anti-α tubulin for microtubules (anti-Tubulin, Santa Cruz Biotechnology, SACSC-53030). The samples were blocked for 1 hour at room temperature and then incubated in secondary antibody solution (Alexa Fluor 488, goat anti-mouse IgG (H+L) Highly Cross-Adsorbed Secondary Antibody, Invitrogen, A-11029, 1:300) overnight at 37°C. For dual color labeling with MAP4-GFP, Alexa Fluor 488 goat anti-rabbit (A11034, Thermo Fisher) and Alexa Fluor 633 goat anti-rat (A21094, Thermo Fisher) secondaries were used. The next day, samples were thoroughly washed in PBS and mounted for inspection in a solution of 90% Glycerol (VWR, ICNA0219399691), 10% PBS, DABCO (25 mg/ml, Sigma Aldrich, D27802), pH 9.

*Multicolor labeling for PlantEx*: We followed a previously published automated labeling protocol^39^. 3 to 4 day-old seedlings were fixed in 24 well-plates with 1 ml of 4% paraformaldehyde in PBS for 1h in the vacuum desiccator with suction applied. Using an Intavis InSitu Pro robot, we incubated 3 times for 15 minutes in PBS + 0.1% Triton X-100 at room temperature (RT), followed by 3 times 15 minutes in ddH_2_O + 0.1% Triton X-100. Initial cell wall digestion was performed for 30 minutes with 2% driselase in PBS at 37 °C. Samples were washed 3 times for 15 minutes in PBS + 0.1% Triton X-100 at RT. Permeabilization was performed for 2 times 30 minutes with 10% DMSO and 3% Igepal CA-630 in PBS at RT, followed by 3 times washing for 15 minutes in PBS + 0.1% Triton X-100 at RT. Blocking was done for 1 h with 2% BSA in PBS. Samples were incubated for 4 h at 37 °C with primary antibodies against GFP, sec21, the γ-subunit of COPI-coated vesicles^62^ (anti-GFP from mouse, monoclonal, Sigma G6539, 1:500; anti-sec21 from rabbit against Arabidopsis thaliana derived sec21-peptide, Agrisera AS08327, 1:500) or non-commercial custom antibodies raised in rabbit against PIN1 or PIN2 in blocking solution and then washed 3 times for 15 minutes at RT in PBS + 0.1% Triton X-100. Incubation with secondary antibodies was done in blocking solution at 37 °C for 4 h (goat anti-rabbit IgG, highly cross adsorbed, conjugated to Alexa Fluor 488, Invitrogen A11034, 1:100; goat anti-mouse IgG, conjugated to Alexa Fluor 594, Abcam, 150116, 1:100). Samples were washed three times for 15 minutes in PBS + 0.1% Triton X-100 at RT and then three times 15 minutes in ddH_2_0 at RT. If anchoring was not done on the same day, samples were stored in storage solution.

*Immunostaining for PlantEx-STED*. We followed an identical protocol as in multicolor labeling above but used a goat anti-rabbit IgG conjugated to Star635P (Abberior, Göttingen, Germany, 1:100).

### Hydrogel embedding and expansion

Whole immunostained seedlings were transferred in PBS onto a large coverslip prepared with a hydrophobic pen (Invignome) to contain fluids. Typically, only one single seedling was placed per coverslip. Further mechanical manipulation or transfers of the seedling were avoided. Fluid was exchanged to anchoring solution (0.2 mg/ml acryloyl-X, SE; Thermo Fisher Scientific, A-20770 in 1x PBS) and the sample was incubated overnight in a humid chamber at RT. For assembly of a gelation chamber similar to the ones previously described^41^, anchoring solution and the hydrophobic rim were removed and a chamber was built up by placing a strip of cover glass on either side of the seedling and covering the assembly with a further 22 mm x 22 mm cover glass on top. The hydrogel monomer solution contained 8.625% (w/w) sodium acrylate (Sigma-Aldrich, 408220), 2.5% (w/w) acrylamide (Sigma-Aldrich, A9099), 0.15% (w/w) N,N’-methylenebisacrylamide (Sigma-Aldrich, M7279), and 2 M NaCl (Sigma-Aldrich, S7653) in PBS. Gelation of monomer solution was initiated by adding 0.2% w/w of a solution containing 10% ammonium persulfate (Sigma-Aldrich, A3678) and 10% TEMED (Sigma-Aldrich, T7024) in ddH_2_O. Gelation chambers were filled with monomer solution and gels were allowed to polymerize for at least 3 hours at 37 °C. After gelation, pre-expansion images were taken on a confocal microscope. For digestion of cell walls, gelation chambers were disassembled with a razorblade and gels were individually placed in wells of a 12-well plate. Digestion proceeded for 24 h at RT in the dark with slight shaking in 2 ml of a solution containing 2% driselase, 1.5% cellulase Onozuka R-10 (Yakult Pharmaceutical Industry Co.), 0.4% macerozyme R-10, 0.2% xylanase (Sigma-Aldrich, X2753) and 0.2% pectolyase Y-23 (Duchefa, P8004) in 1x PBS. Prior to application, the enzyme cocktail was allowed to sediment for 30 minutes and the supernatant was filter sterilized (0.22 μm filter) to remove debris. Gels were then washed twice in PBS. Mechanical homogenization was performed in 2 ml of proteolysis solution containing proteinase K and Ca^2+^ ions (8 U/ml proteinase K (Sigma-Aldrich, P4850), 50 mM TRIS (AppliChem, A3452), 800 mM guanidine HCl (Sigma-Aldrich, G4505), 2 mM CaCl_2_ (Sigma-Aldrich, C5670), 0.5% (v/v) Triton X-100 (Sigma-Aldrich, 93426) in ddH_2_O, pH 8.0) overnight at 50 °C in the dark.

For expansion, gels were placed in 12 cm diameter dishes in the dark with an excess of ddH_2_O. Water was exchanged every 15 minutes until gels were fully expanded, typically after 3 fluid exchanges.

### Pan-protein staining

Hydrogels were incubated with NHS-ester modified dye (40µM in PBS) for 2h at RT with gentle shaking. For confocal measurements, NHS-Alexa488 (APC-002-1, Jena Bioscience, Germany) was used. For STED imaging, NHS-STAR RED (STRED-0002-1MG, Abberior, Germany) was used. After labeling, samples were washed 3x for 15mins in PBS and expanded in ddH_2_O before imaging.

### Proteolysis assay

Protease activity was assessed with the EnzChek Protease Assay Kit (E6638, Thermo Fisher) according to the manufacturer’s instructions. Concentrations of the PlantEx cell wall digestion cocktail and proteinase K, buffers, incubation times, and temperatures were adjusted to the respective experimental steps in the PlantEx protocol. Fluorescence readout was performed with a spectrophotometer. For experiments involving protease inhibitor, one tables of protease inhibitor (cOmplete, 11836153001, Sigma Aldrich) was added to the PlantEx cell wall digestion cocktail. Experiments involving enzymes targeting specific cell wall components were done with α-Amylase 10 U/ml, α-L-Arabinofuranosidase 2.5 U/ml, β-Mannanase 50 U/ml, Cellulase 14 U/ml, Pectate Lyase 5 U/ml, Xyloglucanase 10 U/ml.

### Imaging

*Sample mounting.* Expanded gels were trimmed to contain the root and transferred to a home-made imaging chamber manufactured as previously described^41^.

*Confocal microscopy.* Pre-expansion and PlantEx images were acquired on Zeiss LSM-800 confocal microscopes equipped with plan-apochromat 10x/0.45 NA or 20x/0.8 NA air and plan-apochromat 40x/1.2 NA water immersion objectives. Excitation wavelengths were 488 nm and 561 nm.

*Spinning-disc confocal microscopy.* An Andor Dragonfly microscope based on a Nikon Ti2E inverted microscope, with an Andor Zyla 4.2 Megapixel sCMOS camera, was used for overview imaging, with continuous wave laser excitation at 488 nm, 561 nm and 637 nm as appropriate. Overview images of single plant seedlings were acquired with a 20x air objective (Nikon CFI P-Apo 20x lambda/NA 0.45/WD 4.0 mm). Andor Fusion software was used for data acquisition.

*STED microscopy.* For PlantEx-STED measurements, expanded samples were imaged on an Abberior Instruments Expert Line STED microscope, with 775 nm STED wavelength and ∼1ns long STED pulses at a repetition rate of 40 MHz. Excitation wavelengths were ∼640 nm and ∼560 nm. Nanosecond time gating of fluorescence detection was used throughout STED measurements with a gating window of 8 ns duration. Samples were imaged with a 60x/1.2 NA water immersion objective. Stated power levels refer to the power at the front lens of the objective and bear an estimated uncertainty of 15 %. STED power was distributed to the *xy*-“doughnut” pattern for resolution increase in the focal plane and the *z*-STED pattern for resolution increase along the optical axis according to the lateral/axial STED power ratio stated below with the imaging parameters. Where indicated, Reduction of State transition Cycles (RESCue)-STED^53^ was applied with the parameters stated below, reducing light exposure and photobleaching. At each scan position, fluorescence photon flux was monitored and lasers were turned off in case fluorescence detection failed to reach a lower threshold or it reached an upper threshold, such that signal could be extrapolated to the full pixel dwell time.

### Imaging parameters

*Figure 1, panels d,e*. Pre-expansion: objective lens 40x/1.2 NA; lateral pixel size 100 nm; total imaging *z*-range 60.75 μm; axial step size 750 nm; pixel dwell time 0.35 μs. PlantEx: objective lens 40x/1.2 NA; lateral pixel size 90 nm; total imaging *z*-range 42 μm; axial step size 750 nm; pixel dwell time 0.66 μs.

*Figure 1, panels g,h.* Pre-expansion: objective lens 40x/1.2 NA; lateral pixel size 100nm, total imaging z-range 21µm, axial step size 1µm; pixel dwell time 0.48µs.

PlantEx: objective lens 20x/0.8 NA; lateral pixel size 150nm; total imaging z-range 52.3µm; axial step size 1.09µm; pixel dwell time 0.49µs.

*Figure 2, panel a.* Pre-expansion: objective lens 10x/0.45 NA; lateral scan step (pixel) size 397 nm; total imaging *z*-range 100 μm; axial step size 2 μm; pixel dwell time 0.48 μs.

PlantEx: objective lens 10x/0.45 NA; lateral pixel size 241 nm; total imaging *z*-range 200 μm; axial step size 20 μm; pixel dwell time 0.4 μs with 4 scans per line.

*Figure 2, panel b.* Pre-expansion: objective lens 40x/1.2 NA; lateral scan step (pixel) size 114 nm; total imaging *z*-range 25 μm; axial step size 1 μm; pixel dwell time 0.43 μs.

PlantEx: objective lens 20x/0.8 NA; lateral pixel size 149 nm; single plane; pixel dwell time 1.18 μs.

*Figure 2, panel c.* Pre-expansion: objective lens 40x/1.2 NA; lateral scan step (pixel) size 114 nm; total imaging *z*-range 38 μm; axial step size 1 μm; pixel dwell time 0.43 μs.

PlantEx: objective lens 20x/0.8 NA; lateral pixel size 149 nm; total imaging *z*-range 62 μm; axial step size 2 μm; pixel dwell time 0.51 μs.

*Figure 3, panel a.* Pre-expansion: objective lens 40x/1.2 NA; lateral scan step (pixel) size 100 nm; total imaging *z*-range 18 μm; axial step size 1 μm; pixel dwell time 0.48 μs.

PlantEx: objective lens 40x/1.2 NA; lateral pixel size 156 nm; total imaging *z*-range 40 μm; axial step size 1 μm; pixel dwell time 0.38 μs.

*Figure 3, panel b*. Pre-expansion: objective lens 40x/1.2 NA; lateral pixel size 100 nm; total imaging *z*-range 51 μm; axial step size 1.5 μm; pixel dwell time 0.48 μs.

PlantEx: objective lens 40x/1.2 NA; lateral pixel size 81 nm; total imaging *z*-range 41 μm; axial step size 1 μm; pixel dwell time 0.4 μs with 4 scans per line.

*Figure 4, panel a, b*.

PlantEx: confocal image; objective lens 60x/1.2 NA; pixel size 30 nm x 30 nm; pinhole size 0.83 Airy Unit; Excitation laser power: 7.2 μW; pixel dwell time 20 μs. PlantEx-STED: objective lens 60x/1.2 NA; pixel size 30 nm x 30 nm; pinhole size 0.83 AU; Excitation laser power: 7.2 μW; STED laser power: 32.5 mW; lateral/axial STED power ratio 100/0; pixel dwell time 63 μs; RESCue STED lower thresholds 1,2,5, and 10 counts after 21%, 28%, 52%, and 85% of total pixel dwell time, respectively; RESCue STED upper threshold 28 counts.

*Figure 4, panel e.* PlantEx: confocal image; objective lens 60x/1.2 NA; pixel size 100 nm x 100 nm; pinhole size 0.83 AU; Excitation laser power: 7.2 μW; pixel dwell time 20 μs. PlantEx-STED: objective lens 60x/1.2 NA; pixel size 100 nm x 100 nm; pinhole size 0.83 AU; Excitation laser power: 7.2 μW; STED laser power: 38 mW; lateral/axial STED power ratio 0/100; pixel dwell time 63 μs; RESCue STED lower thresholds 1,2,5, and 10 counts after 21%, 28%, 52%, and 85% of total pixel dwell time, respectively; RESCue STED upper threshold 28 counts.

*Figure 4, panel g,h,i.* PlantEx-STED: objective lens 60x/1.2 NA; voxel size 100 nm x 100 nm x 100 nm; pinhole size 0.83 AU; Excitation laser power: 7.2 μW; STED laser power: 21.7 mW; lateral/axial STED power ratio 0/100; pixel dwell time 63 μs; RESCue STED lower thresholds 1,2,5, and 10 counts after 21%, 28%, 52%, and 85% of total pixel dwell time, respectively; RESCue STED upper threshold 28 counts.

*Figure 4, panel j.* PlantEx-STED: objective lens 60x/1.2 NA; voxel size 100 nm x 100 nm x 100 nm; pinhole size 1 AU; Excitation laser power: 14.2 μW; STED laser power: 30 mW; lateral/axial STED power ratio 0/100; pixel dwell time 60 μs; volume size 10x10x80µm; *xyz* scan mode; pinhole 1 airy unit; RESCue STED lower thresholds 1,2,5, and 10 counts after 21%, 28%, 52%, and 85% of total pixel dwell time, respectively; RESCue STED upper threshold 28 counts.

*Figure 5, panel a.* Spinning disc confocal image (Andor Dragonfly), objective lens 20x/0.75 NA; pinhole size 25µm; lateral pixel size 300 nm; total imaging *z*-range 73 μm; axial step size 3 μm; 488nm laser power: 14%; exposure time 100 ms, Andor Zyla sCMOS camera. Software: Andor Fusion version 2.2.0.49.

*Figure 5, panel b.* PlantEx-STED: objective lens 60x/1.2 NA; pixel size 30 nm x 30 nm; pinhole size 1 AU; Excitation laser power 488nm: 1.9 µW; 560nm: 3.3 µW; 640nm: 14.2 µW; STED laser power: 10 mW; lateral/axial STED power ratio 100/0; pixel dwell time 60 μs.

*Figure 5, panel c.* PlantEx-STED: objective lens 60x/1.2 NA; pixel size 30 nm x 30 nm; pinhole size 1 AU; Excitation laser power 488nm: 5.6 μW; 560nm: 6.6 µW; 640nm: 14.2 µW; STED laser power: 10 mW; lateral/axial STED power ratio 100/0; pixel dwell time 40 μs.

*Figure 5, panel d.* PlantEx-STED: objective lens 60x/1.2 NA; pixel size 30 nm x 30 nm; pinhole size 1 AU; Excitation laser power 560nm: 5.3 µW; 640nm: 1.4 µW; STED laser power: 5 mW; lateral/axial STED power ratio 100/0; pixel dwell time 50 μs.

*Figure 5, panel e.* PlantEx-STED: objective lens 60x/1.2 NA; voxel size 100 nm x 100 nm x 100 nm; pinhole size 1 AU; Excitation laser power: 14.2 μW; STED laser power: 30 mW; lateral/axial STED power ratio 0/100; pixel dwell time 60 μs; Scan mode *xzy*; RESCue STED lower thresholds 1,2,5, and 10 counts after 21%, 28%, 52%, and 85% of total pixel dwell time, respectively; RESCue STED upper threshold 28 counts.

*Suppl.* Figure 3*, panels a,b.* Pre-expansion: objective lens 40x/1.2 NA; lateral pixel size 100nm, total imaging z-range 8.4µm, axial step size 0.3µm; pixel dwell time 0.59µs. PlantEx: dataset 1: objective lens 40x/1.2 NA; lateral pixel size 100nm; total imaging z-range 23µm; axial step size 1µm; pixel dwell time 1.31µs. PlantEx: dataset 2: objective lens 40x/1.2 NA; lateral pixel size 100nm; total imaging z-range 34µm; axial step size 1µm; pixel dwell time 1.31µs.

*Suppl.* Figure 4, PlantEx: confocal image; lateral pixel size 100 nm x 100 nm; pinhole size 0.6 AU; Excitation laser power: 21.6 μW; pixel dwell time 20 μs. PlantEx-STED: pixel size 25 nm x 25 nm; pinhole size 0.6 AU; Excitation laser power: 21.6 μW; STED laser power: 21.7 mW; lateral/axial STED power ratio 100/0; pixel dwell time 20 μs.

### Image analysis

*Processing of images.* Images represent raw confocal or STED imaging data. No deconvolution or image restoration was applied. Intensity lookup tables, maximum intensity projections and color-coded projections were applied with FIJI/ImageJ, version 1.51p, 1.52p, 1.54f or 1.53t. For further image processing and for clustering of vesicles, we used Python 3.6-3.10 together with the scikit-image^63^ (ver. 0.20.0) library.

*Image registration, determination of expansion factor, and distortion analysis.* Distortion analysis was adapted from previously published methods^44^ (https://github.com/Yujie-S/Click-ExM_data_process_and_example/tree/master). Pre- and post-expansion volumes were imaged with a confocal microscope, allowing quantification of distortions above the diffraction limit (∼200 nm). The BigWarp^64^ Fiji-plugin (v. 7.0.7) was used for registration of corresponding pre- and post-expansion imaging volumes. First, we placed ∼20 landmarks at corresponding plant features in pre- and post-expansion imaging volumes. We then applied a similarity transformation (isotropic scaling, translation, and rotation) to the pre-expansion volume to overlap landmarks and align to the fixed post-expansion volume and crop to the same axial extent of the volumes. The expansion factor was extracted as the linear scaling factor of the similarity transformation minimizing squared landmark residuals using the script https://github.com/danzllab/CATS/tree/master/rcats_image-analysis/bigwarp. For Fig. 2b, we placed landmarks in 2D, as the post-expansion image was recorded as a single optical slice. For scaling, we either used the expansion factor determined for the specific experiment or the average expansion factor determined from *n*=12 specimens (Fig. 1c). We then created maximum intensity projections from the aligned pre-/post-expanded volumes, selecting axial ranges for intensity projects to cover similar ranges in the sample, as judged by visual inspection. To make resolution and appearance of pre- and post-expansion images similar, we smoothed them with 2D Gaussian filters of different sizes (σ=1 and σ=6 pixels, respectively).

For the distortion analysis in Figure 2a, we separated foreground signal from background: First, the preprocessed pre- and post-expansion projections were background-subtracted using a strong 2D Gaussian blur (σ = 80). Resulting negative values were clipped to zero. Then, we created signal-containing foreground masks by thresholding. A mask from pre- or post-expansion images was either applied directly or they were combined into a single binary mask by a logical OR operation. Finally, we applied morphological closing operation to remove small, spurious holes in the mask. Thresholds and sizes of the closing operation were chosen by visual inspection.

To quantify non-linearities in the expansion procedure, we computed distortion vector fields from the pre- and post-expansion images using the *imregdemons.m* function in MATLAB (version R2022b, MathWorks). We calculated the measurement error across different measurement lengths, as previously described^19^. For this, we randomly sampled 200,000 pairs of feature points from the segmented foreground. The distortion field is applied to each pair to retrieve its corresponding transformed vector. We then calculated the measurement errors for different measurement lengths by subtracting corresponding feature vectors and taking the length of the difference vectors. For Figures 2e, the root mean square (RMS) of measurement errors was calculated across the *n*=4 specimens in Fig. 2a-d. For all figures, the RMS errors, measurement errors, and measurement lengths are shown in the pre-expansion scale.

*Line profiles*. Line profiles were created with FIJI/ImageJ, version 1.51n and 1.54f. Line profiles were created from single confocal or STED image planes. For noise reduction, averaging of 3 pixel width perpendicular to the direction of the line profile was applied.

*Vesicle FWHM quantification in 2D.* In the raw PlantEx-STED images, we first normalized the raw photon counts from 16-bit unsigned integer values to floating point values by dividing by the 99.9 percentile of the raw brightness values. We then applied an isotropic 2D Gaussian filter with a sigma of size 3.3 pixel (100 nm in expanded sample) followed by the detection of local maxima in 2D. To suppress noisy background detections, we filtered for maxima having a brightness value greater than 25% of the maximum value in the image. We then selected isolated vesicles that were separated by at least 30 pixels (900nm in expanded sample) from other detections. We fitted an isotropic Gaussian function to these isolated vesicles. For the fitting, we extracted a square sub-image of 41-pixel width (1.23 µm after expansion) centered at each isolated vesicle from the raw data. In each sub-image, we fitted a Gaussian with 4 degrees of freedom (amplitude, isotropic standard deviation, center sub-pixel position in two spatial dimensions) or 5 degrees of freedom (amplitude, standard deviation in two directions, center sub-pixel position in two spatial dimensions) by minimizing squared residuals using the Levenberg-Marquardt algorithm. Upon convergence, we calculated the full-width-at-half-maximum (FWHM) by multiplying the standard deviation with 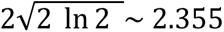 and used the measured expansion factor to convert to the native tissue scale. We manually filtered the vesicles where the 2D Gaussian fitting failed due to background noise, overlap of neighboring peaks or signal from out-of-focus planes. In total *n*=74 vesicles across *n*=3 samples were analyzed. Gaussian fitting was done using the scipy library (ver. 1.12.0). We implemented the described routine as Python command line script *gaussian_fit_fwhm* for 2D and 3D image data. For the vesicle FWHM estimate, we calculated a median value with its 25th and 75th percentile.

*Vesicle FWHM quantification in 3D*. In the raw volumetric PlantEx-STED images, we first normalized the raw photon counts from 16-bit unsigned integer values to floating point values by dividing by the 99.9 percentile of the raw brightness values. We then applied an isotropic 3D Gaussian filter with a sigma of 1 voxel (100 nm in expanded sample) followed by the detection of local maxima in 3D. To suppress noisy background detections, we filtered for local maxima having a brightness value greater than 25% of the maximum value in the image. We then selected isolated vesicles that were separated by at least 18 voxels (1.8 µm after expansion) from other detections and fitted an isotropic 3D Gaussian function to these. For the fitting, we extracted cubes of 21 voxel edge length (2.1 µm after expansion) centered at each isolated vesicle from the raw data. In each sub-volume, we fitted a 3D Gaussian with 5 degrees of freedom (amplitude, isotropic standard deviation and center sub-pixel positions in three spatial dimensions) or 7 degrees of freedom (amplitude, standard deviation in three directions, center sub-pixel position in three spatial dimensions) by minimizing squared residuals using the Levenberg-Marquardt algorithm. Upon convergence, we calculated the full-width-at-half-maximum (FWHM) and used the measured expansion factor to convert to the native tissue scale. In total, *n*=112 vesicles were analyzed across *n*=3 specimens.

*Identification and visualisation of 3D clusters.* In the raw PlantEx-STED images, we first normalized the raw photon counts from 16-bit unsigned integer values to floating point values by dividing by the 99.9 percentile of the raw brightness values. We then applied an isotropic Gaussian filter with a sigma of 0.7 pixel (70 nm after expansion), followed by the detection of local maxima in 3D. The minimum allowed separation between any two local maxima was set to 2 voxel edge lengths (200nm). If two local maxima were closer, we chose the maximum with the higher brightness value. To suppress noisy background detections, we filtered for maxima with a brightness of at least 10% of the maximum value in the image. In order to identify clusters of sec21-positive puncta, we applied the OPTICS algorithm^65^. We set the input parameter for the minimum cluster size to 10 (minimum number of vesicles to form a cluster). All other parameters were left to their default values (scikit-learn ver. 0.21.3). 3D-rendering of vesicles was done using the SciView library^66^ in FIJI/ImageJ, version 1.51p.

