## Supplementary Information for "Super-resolution expansion microscopy in plant roots"

[1]Institute of Science and Technology Austria, Am Campus 1, 3400 Klosterneuburg, Austria
[2]present address: Medical University of Vienna, Division of Anatomy, Centre for Anatomy & Cell Biology, Vienna, Austria
[3]present address: University of Exeter, Biosciences, Exeter, UK

†authors contributed equally

Short title: Expansion microscopy in plants

### Contents

**Supplementary Figure 1: Non-specific proteolytic activity of cell wall digestion cocktail** 2

**Supplementary Figure 2: Alignment of samples for distortion analysis** 3

**Supplementary Figure 3: Mitotic figures in tubulin-labeled PlantEx samples** 4

**Supplementary Figure 4: STED imaging in *A. thaliana* root without PlantEx** 5

**Supplementary Figure 5: 3D-rendering of COPI-coated vesicle distribution imaged with PlantEx-STED** 6

**Supplementary Figure 6: FWHM of COPI-coated vesicles in PlantEx-STED.** 7

**Supplementary Figure 7: PlantEx pan-protein labeling with non-inverted intensity lookup table.** 8

**Captions for Supplementary Movies.** 9

**Supplementary Figure 1: Non-specific proteolytic activity of cell wall digestion cocktail**

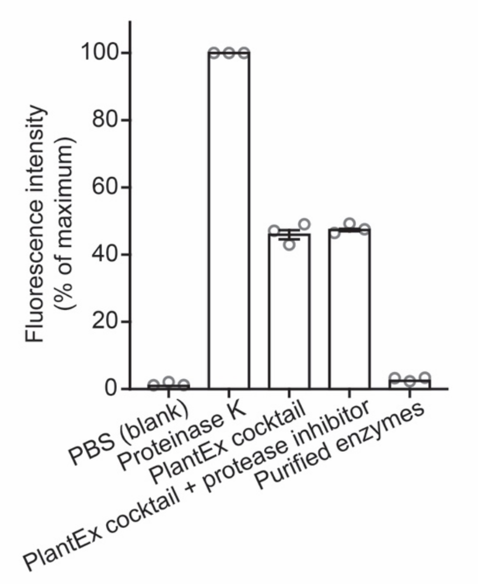


**Non-specific proteolytic activity of cell wall digestion cocktail.** Fluorescence assay for non-specific proteolytic activity (EnzChek, Thermo Fisher), with fluorescence readout intensity normalized to proteinase K activity. Concentrations, incubation times, and temperatures for proteinase K and PlantEx cell wall digestion cocktail were chosen identical to the respective digestion steps in PlantEx experiments. The PlantEx cell wall digestion cocktail contains natural products that are not chemically defined (fungal extracts) and displayed 46 ± 3 % (mean ± s.d.) of non-specific proteolytic activity relative to proteinase K. Proteolytic activity was not suppressed by a proprietary protease inhibitor mix (PlantEx cocktail + protease inhibitor, cOmplete, Sigma Aldrich). A mixture of purified enzymes targeting specific cell wall components ($\alpha$-amylase, $\alpha$-L-arabinofuranosidase, $\beta$-mannanase, cellulase, pectate lyase, xyloglucanase) we tested for PlantEx did not display substantial proteolytic activity but was not effective at achieving mechanical homogenization for expansion. Mean ± s.d., datapoints represent 3 experimental repetitions.

**Supplementary Figure 2: Alignment of samples for distortion analysis**

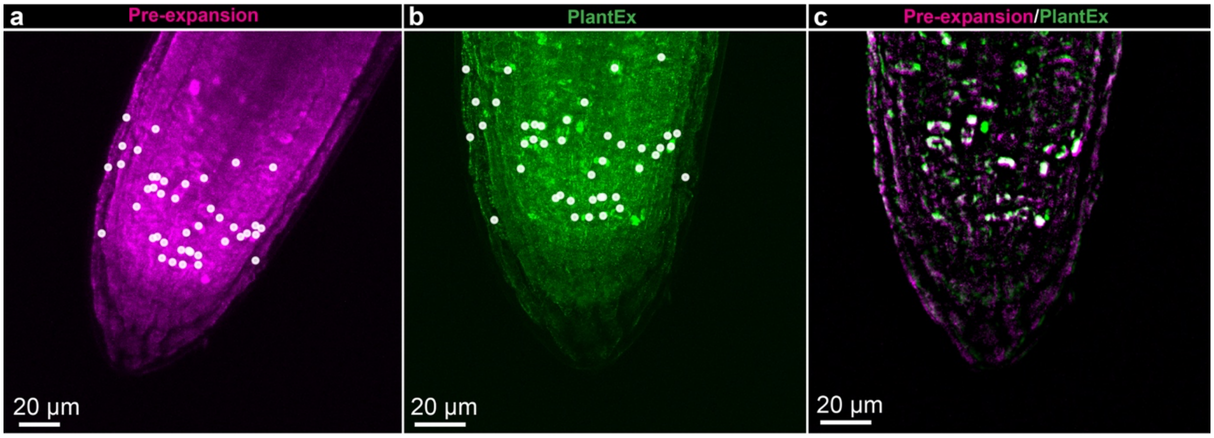


**Alignment of samples for distortion analysis**. **a,b,** Manually placed landmarks (white) overlaid with a maximum intensity projection of the pre- and post-expansion confocal imaging stacks in **Fig. 2a**. For alignment, landmark features that were readily identified in both pre-expansion and PlantEx image stacks were marked using the BigWarp plugin for ImageJ. Note that the landmarks were not actually set on the shown maximum intensity projections, but on individual slices of the image stack, to maximize alignment fidelity. For alignment of pre-expansion and PlantEx images, a (linear) similarity transformation with the following 7 degrees of freedom was performed: isotropic scaling (equivalent to expansion factor), rotation (3 angles), and 3D-translation (3 axes). Scale bars: 20 µm (corresponding to 82 µm in expanded sample, exF=4.1). **c**, Overlay of pre- and post-expansion images. The homogeneous signal component was removed by Gaussian background subtraction to focus distortion analysis on distinct image features.

### Supplementary Figure 3: Mitotic figures in tubulin-labeled PlantEx samples

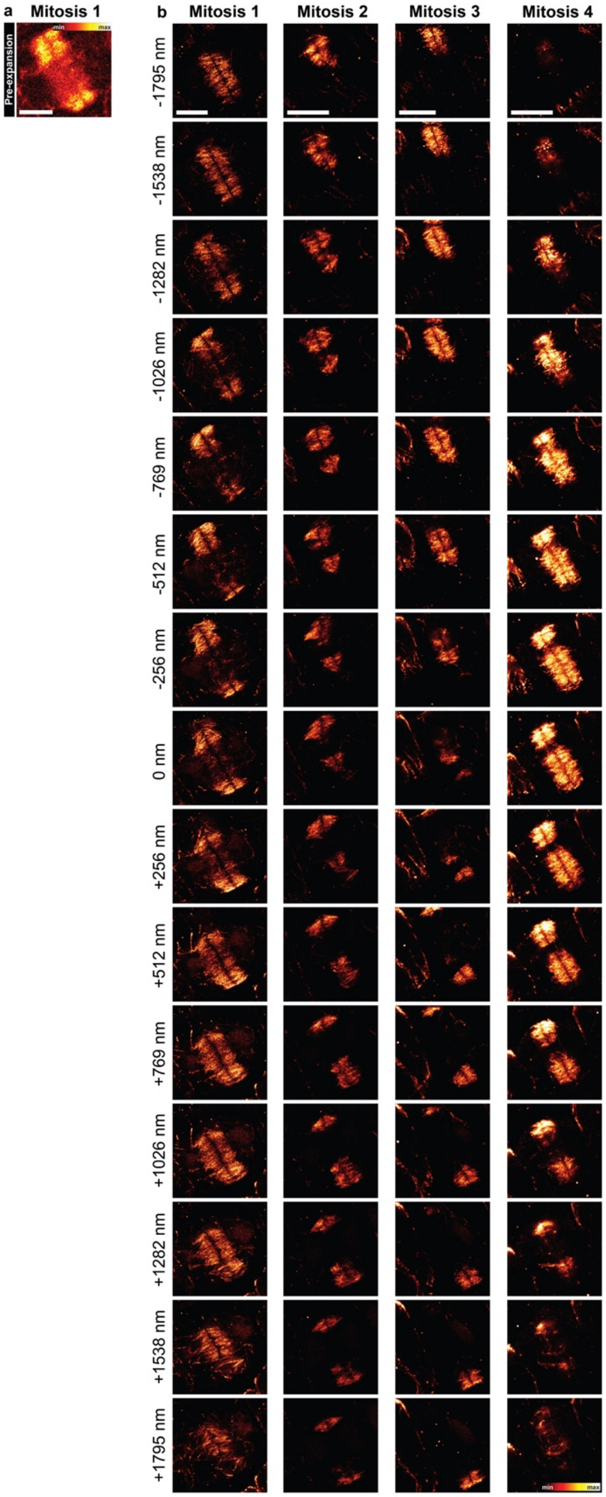


**Mitotic figures in tubulin-labeled PlantEx samples**. **a,** Confocal image from a non-expanded *A. thaliana* root sample immuno-labeled for tubulin. Sample was treated with taxol prior to fixation to increase number of mitotic arrests. **b,** Confocal image stacks in the same root after application of PlantEx, showing the same mitotic figure at 4-fold increased resolution (mitosis 1) and 3 further examples from the same specimen. Scale bars: 5 µm.

**Supplementary Figure 4: STED imaging in *A. thaliana* root without PlantEx**

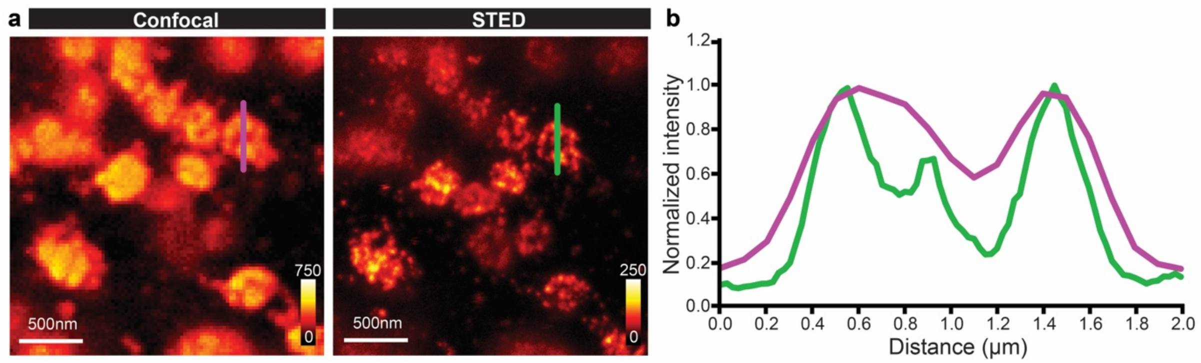


**STED imaging in *A. thaliana* root without PlantEx. a,** *A. thaliana* root tip immuno-labeled for Sec21 and imaged with confocal and STED microscopy (with lateral resolution increase, *xy*-STED pattern), respectively. No hydrogel expansion was applied. Scale bar: 500 nm. **b,** Line profiles as indicated in panel a. *A. thaliana* roots are challenging samples for STED imaging, due to factors including scattering and refractive index mismatch and variation.

**Supplementary Figure 5: 3D-rendering of COPI-coated vesicle distribution imaged with PlantEx-STED**

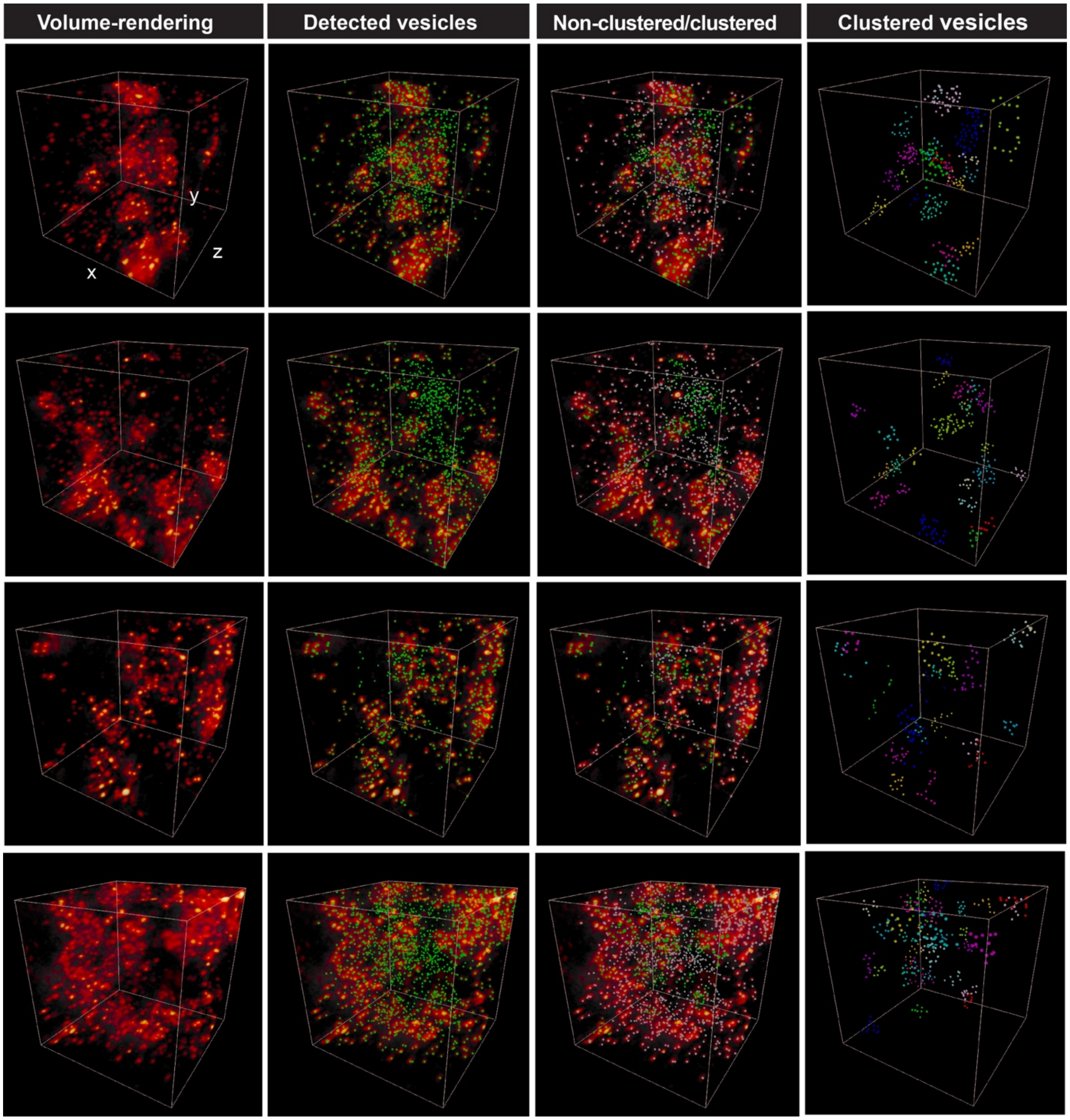


**3D-rendering of COPI-coated vesicle distribution imaged with PlantEx-STED.** Volume renderings from a PlantEx *Arabidopsis thaliana* whole-mount root sample immunostained for COPI-coated vesicles (γ-COP) and imaged with near-isotropic STED microscopy (*z*-STED pattern to increase resolution predominantly in the axial direction). Imaging volumes were 15 x 15 x 15 $\mu m^3$ after expansion, corresponding to 3.8 x 3.8 x 3.8 $\mu m^3$ in the native tissue (column 1). For visualization, γ-COP positive puncta were detected as local intensity maxima (column 2) and classified as clustered (green) or non-clustered (grey) by the OPTICS algorithm[1] (column 3), with a minimum cluster size of 10. The last column shows only detections assigned to clusters, with individual clusters color coded. Imaging volumes contained 925, 1066, 548, and 1497 detections in the 4 imaging volumes from top to bottom.

**Supplementary Figure 6: FWHM of COPI-coated vesicles in PlantEx-STED.**

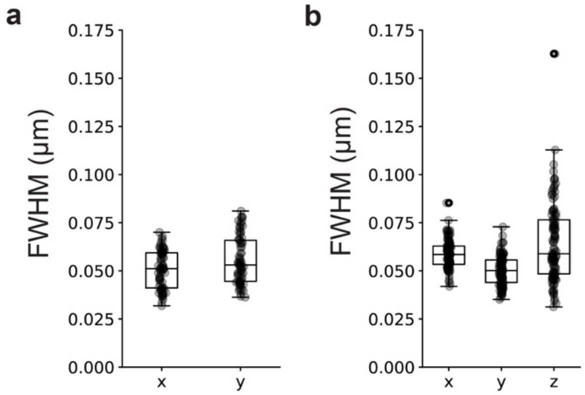


**FWHM of COPI-coated vesicles in PlantEx-STED**. **a,** Anisotropic 2D Gaussian fits (5 degrees of freedom: amplitude, width in 2 directions, position in 2D) of the same data as in Fig. 4d. FWHM median (lower, upper quartiles) of the 74 vesicles analyzed across *n*=3 specimens: *x*: 51 nm (41 nm, 59 nm); *y*: 53 nm (45 nm, 66 nm). **b,** Anisotropic 3D Gaussian fits (7 degrees of freedom: amplitude, width in 3 directions, position in 3D) to the same data as in Fig. 4f. FWHM median (lower, upper quartile) of the 112 vesicles analyzed across *n*=3 specimens: *x*: 59 nm (53 nm, 63 nm); *y*: 50 nm (44 nm, 56 nm); *z*: 59 nm (48 nm, 77 nm). Here, application of a *z*-STED pattern with resolution increase predominantly in the axial direction leads to near-isotropic overall resolution.

**Supplementary Figure 7: PlantEx pan-protein labeling with non-inverted intensity lookup table.**

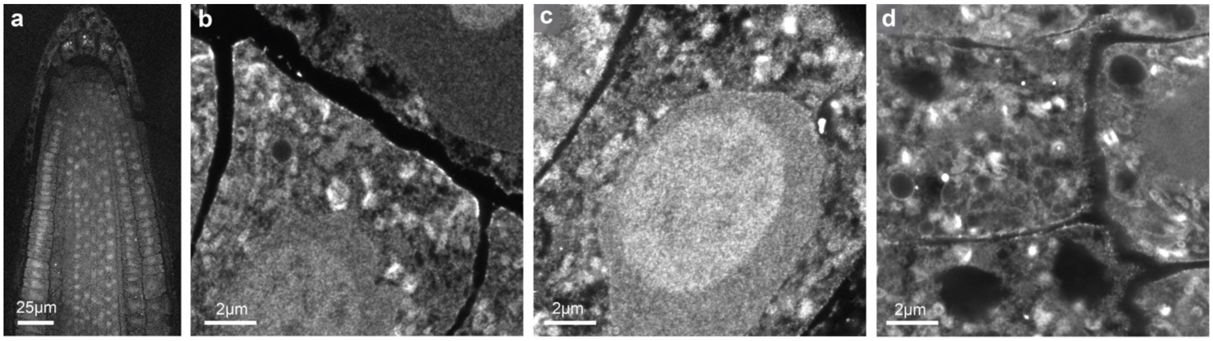


**PlantEX pan-protein labeling with non-inverted color map.** Same data as in Fig. 4a-d, without inverting the intensity lookup table. Black: low protein density. White: high protein density.

### Captions for Supplementary Movies.

*Suppl. Movie 1.* COPI-coated vesicles (green) and trans-Golgi network (magenta) in conventional confocal microscopy before expansion. Image stack of a larger region from the same sample as in Fig. 2d.

*Suppl. Movie 2.* COPI-coated vesicles (green) and trans-Golgi network (magenta) with PlantEx, imaged with confocal microscopy. Image stack of a larger region from the same sample as in Fig. 2f.

*Suppl. Movies 3-6*. COPI-coated vesicles in tissue volumes imaged with PlantEx-STED. Fly-through of the raw imaging data along the optical axis for individual tissue volumes displayed in Fig. 4g and Suppl. Fig. 5. Scale bars refer to original tissue scale.

*Suppl. Movie 7-10*. Movies of the 3D renderings of COPI-coated vesicles imaged with PlantEx-STED at near-isotropic resolution in the tissue volumes in Fig. 4g and Suppl. Fig. 5. First the imaging data is shown, then in addition vesicle detections. Next, they are classified into vesicles assigned to clusters (green) and those outside clusters (gray) and subsequently only the clustered vesicles are shown. Clusters are then color coded.
